# Human and Rodent Temporal Lobe Epilepsy is Characterized by Changes in O-GLCNAC Homeostasis that can be Reversed to Dampen Epileptiform Activity

**DOI:** 10.1101/330738

**Authors:** Richard G. Sanchez, R. Ryley Parrish, Megan Rich, William M. Webb, Roxanne M. Lockhart, Kazuhito Nakao, Lara Ianov, Susan C. Buckingham, Mark Cunningham, Devin R. Broadwater, Alistar Jenkins, Nihal C De Lanerolle, Tore Eid, Kristen Riley, Farah D. Lubin

## Abstract

Temporal Lobe Epilepsy (TLE) is frequently associated with changes in protein composition and post-translational modifications (PTM) that exacerbate the disorder. O-linked-β-N-acetyl glucosamine (O-GlcNAc) is a PTM occurring at serine/threonine residues that integrate energy supply with demand. The enzymes O-GlcNActransferase (OGT) and O-GlcNAcase (OGA) mediate the addition and removal, respectively, of the O-GlcNAc modification. The goal of this study was to determine whether changes in OGT/OGA cycling and disruptions in protein O-GlcNAcylation occur in the epileptic hippocampus. We observed reduced global and protein specific O-GlcNAcylation and OGT expression in the kainate rat model of TLE and in human TLE hippocampal tissue. Inhibiting OGA with Thiamet-G elevated protein O-GlcNAcylation, and decreased both seizure duration and epileptic spike events, suggesting that OGA may be a therapeutic target for seizure control. These findings suggest that loss of O-GlcNAc homeostasis in the kainate model and in human TLE can be reversed via targeting of O-GlcNAc related pathways.

## 1 Introduction

Temporal lobe epilepsy (TLE) is a neurological disorder characterized by recurrent, unprovoked seizures. Previous studies in TLE have revealed changes in cytoskeleton modifications, synaptic proteins, mitochondrial proteins, ion channels, and chaperone proteins [1, 2]. Although proteomic studies have investigated the role of post-translational modifications (PTM) in these proteins, these studies have focused mainly on phosphorylation. While associations between O-GlcNAcylation and hyper-excitability have been reported, this study examines the relationship between protein O-GlcNAcylation and epilepsy and considers O-GlcNAc-related pathways as a potential therapeutic target in epilepsy [3, 4].

Protein expression and function is a dynamic process that requires precise regulation in order to maintain cellular homeostasis under changing conditions. One process by which cells regulate these parameters is PTMs, where enzymes add functional groups to modulate protein dynamics. In TLE, several of these modifications are disrupted, with the majority of studies to date revealing irregularities in the extent of protein phosphorylation and its downstream effects on neuronal homeostasis [5-7]. Because of its similarities to phosphorylation, O-GlcNAcylation, has recently gained attention in hyper-excitability studies [8-10]. O-GlcNAcylation depends on cellular metabolism in order to synthesize the substrate, UDP-GlcNAc, which is then used by O-GlcNAc Transferase (OGT) to add O-GlcNAc to serine and threonine residues [8-10]. The removal of this modification is regulated by O-GlcNAcase (OGA) [8-10]. Together, OGT and OGA regulate global levels of O-GlcNAcylation across a variety of cellular stresses in order to preserve homeostasis [11]. Unlike phosphorylation, in which numerous kinases and phosphatases target many of the same proteins, O-GlcNAcylation relies on only OGT and OGA [12]. Importantly, OGA can be selectively inhibited by Thiamet-G, a purine analog that can cross the blood-brain barrier [13]. By inhibiting OGA, Thiamet-G can thus be used to increase global levels of O-GlcNAcylation within eukaryotic cells [13]. Recently, several studies have focused on O-GlcNAcylation in neurological disorders such as Alzheimer’s, Parkinson’s, Huntington’s, Schizophrenia, and other nervous system processes such as appetite, LTD, hyper-excitability and protein structure [14-23]. However, the role of OGT/OGA cycling and protein O-GlcNAcylation in the epileptic hippocampus and the question of whether O-GlcNAc pathways could be targeted for treatment of TLE remains to be determined.

In the present study, we investigated the role of O-GlcNAcylation both in a rodent model of TLE and also in human epileptic brain tissue, asking whether targeted manipulation of this modification could ameliorate epileptiform brain activity. We identified global decreases in O-GlcNAcylation in epileptic rats and in human patients with TLE. Mass spectrometry studies revealed that O-GlcNAcylation marks were irregularly expressed on specific proteins in the hippocampi of epileptic rats compared to age-matched non-epileptic controls.

Furthermore, we found that inhibition of OGA in rodents using Thiamet-G resulted in improved seizure behavior and decreased interictal spike frequency. Similarly, electrophysiological recordings from human TLE samples showed decreased spike events with OGA inhibition compared to recordings taken in vehicle-treated controls. Collectively, these results support a critical role for protein O-GlcNAcylation in epilepsy and its novel therapeutic potential in the treatment of chronic seizures.

## 2 Material and Methods

### 2.1 Antibodies

The following antibodies were used: 1:500 anti-O-GlcNAc (CTD110.6,-MMS-248R from Covance, Princeton, NJ, USA), 1:500 anti-O-GlcNAc Transferase (O6264, Sigma, St. Louis, MO, USA), 1:20000 goat-anti-mouse (926-32350, Licor, Lincoln, NE, USA), 1:20000 goat-anti-rabbit (926-32211, Licor), 1:1000 anti-Actin (ab1801, Abcam, Cambridge, UK), 1:1000 anti-NeuN (MAB377, Abcam), 1:1000 anti-GFAP (ab7260, Abcam).

### 2.2 Electroencephalogram (EEG)

4 weeks following the administration of kainic acid, rats underwent an electrode implantation. Electrodes (MS333/1-B/SPC, Plastics One, Ranoke, VA, USA) for EEG recordings were trimmed to 1.75 mm in length and fitted into three holes so that they contacted the dura and the connector was flush with the skull. The ground wire was placed into the most caudal hole. For EEG recordings, animals were transferred to individual housing in custom-designed and constructed plexiglass cages at 5 weeks. EEG data were acquired using 8 Biopac Systems amplifiers and AcqKnowlege 4.1 EEG Acquisition and Reader Software (BIOPAC Systems, Inc., Goleta, CA, USA). Data were stored and analyzed in digital format. Each cage was also equipped with an IR Digital Color CCD camera (Lorex Technology, Inc., Linthicum, MD, USA) and animals are recorded concurrently with EEG monitoring. Baseline recordings were done for 24hrs then 10mg/kg of Thiamet-G dissolved in 0.1% w/v saline (SD Chemmolecules, Owings Mills, MD, USA) was administered intraperitoneal (I.P) and then 10mg/kg after each post-injection. Both saline and kainic acid, cohorts were injected with Thiamet-G. After 24 hrs of EEG recording post-injection, a second treatment with the same dosage was administered. Animals were recorded via EEG for 24 hrs after each injection of Thiamet-G. Animals received a total of three independent treatments at same dosage of the drug.

Tissue for Western blots was collected from 4 weeks of age using the previously-described methods. The whole hippocampus was collected and then sub-dissected. All EEG data were analyzed manually using Matlab by an observer blinded to the sample’s identity. Abnormalities in the recordings indicative of epileptic activity are aligned chronologically with the corresponding video in order to confirm seizures.

### 2.3 Immunofluorescence

Animals were sacrificed by rapid decapitation; brains were removed, and fixed in 4% paraformaldehyde overnight at 4°C. The next day the samples were washed with 1x PBS 5x five minutes each time before incubating with 30% sucrose (w/v) overnight at 4°C. The tissue was then flash frozen on dry ice and mounted in O.C.T. (VWR, Randor, PA, USA) 10-micron Sections (10µM) were taken throughout the dorsal hippocampus and mounted onto slides. Antigen retrieval was done by boiling in citric acid buffer followed by washing in 1x PBS. Slices were then blocked for 1hr (4% normal goat serum, 4% normal donkey serum and 0.3% Triton-X in PBS) and incubated in primary antibody for O-GlcNAc (1:200 CTD110.6, MMS-248R, Covance), NeuN (1:1000, MAB377, Millipore), and GFAP (1: 1000, ab7260, Abcam), overnight at 4°C. The following day sections were rinsed with 1x PBS and incubated in Alexa Fluor 488-labeled (1:500, #111-545-003, Jackson Immuno Research, West Grove, PA, USA) or Rhodamine-labeled (TRITC, 1:500, #715-025-150, Jackson Immuno Research) secondary antibodies for 2hrs and, rinsed with 1x PBS and then coverslipped with Vectashield mounting media with DAPI (H-1500, Vector Laboratories, Burlingame, CA, USA). Images were taken on a Zeiss Axio Imager microscope and analyzed using Image J.

### 2.4 Human Tissue Samples

Pharmacologically-resistant hippocampal and cortical tissue samples from human TLE patients were provided by Tore Eid, MD from the Departments of Laboratory Medicine and of Neurosurgery, at Yale School of Medicine. Additional tissue was provided by Kristen O. Riley, MD from the Department of Neurology and Yancy G. Gillespie, MD from the Wallace Tumor Institute at UAB. Acquisition and processing of control human tissue were performed by the Alabama brain collection https://www.uab.edu/medicine/psychiatry/research/resources-0/alabama-brain-collection. Patient demographics and pharmacological history are described in table 2.

**Table 1:** Differentially expressed proteins and their O-GlcNAcylation levels in epilepsy.

| Protein | NP# | Role | O-GlcNAcylation | References |
| --- | --- | --- | --- | --- |
| Acyl-CoA synthetase family member 2, mitochondrial | NP_001030125.1 | Catalyzes the initial reaction in fatty acid metabolism; thioester with CoA | ↑ |  |
| Catenin, alpha 1 | NP_001007146.1 (+1) | RNA binding structural molecule | ↓ |  |
| Mitochondrial import receptor subunit TOM34 | NP_001037709.1 | Aids in imports from cytosol to mitochondria | ↓ |  |
| Filamin C | NP_001178791.1 (+1) | Actin-cross-linking protein | ↓ | Lee et al, Petrecca et al |
| Filamin A | NP_001128071.1 (+1) | Actin binding protein | = | Parini et al, Sheen et al, Hehr et al |
| Fused in sarcoma RNA-binding protein | NP_001012137.1 (+1) | Binds to RNA, and interacts with nuclear pore and transcription initiation factors | ↓ | Neuman et al, Wolfe et al |
| Afadin | NP_037349.1 | Ras target that regulates cell-cell adhesions | ↓ | Yamamoto et al |
| ATP synthase subunit alpha, mitochondrial | NP_015581.1 | Produces ATP | = | Zsoka et al |
| Cathepsin D | NP_099161.2 | Lysosomal aspartyl protease | ↑ | Herman et al, Zhao et al |
| CCR4-NOT transcription complex, subunit 1 | NP_001128312.1 | Part of the CCR4-NOT complex that deadenylates mRNA's | ↑ |  |
| EH- Domain containing protein 3 | NP_620245.2 | Membrane tubulation/endocytic transport | ↓ |  |
| Sorulin-related receptor precursor | NP_445971.1 | Endocytic receptor, possibly for lipoproteins, and proteins | ↑ | VonDemn et al, Friedman et al, Volosin et al, Hiveron et al |
| Dual specificity mitogen-activated protein kinase kinase 4 | NP_001025194.1 | MAP kinase | ↓ | Choi et al, Waltereit et al, Salzman et al |
| Glycerol-3-phosphate dehydrogenase, mitochondrial | NP_036868.1 | Enzyme that plays a role in lipid biosynthesis | ↑ | Eun et al, Link et al |
| A-kinase anchor protein 5 | NP_398199.2 | May anchor PKA to the cytoskeleton/organelles | ↑ | Kay et al, Zhang et al, Tunquist et al |
| Protein kinase C substrate 80K-H | NP_001100276.1 (+2) | Regulatory subunit of glucosidase II | ↓ | De Prater et al, Dardin et al |
| Ras-specific guanine nucleotide-releasing factor 2 | NP_446173.1 | Guanine nucleotide releasing factor for RAS | ↑ | Zhu et al |

**Table 2:**
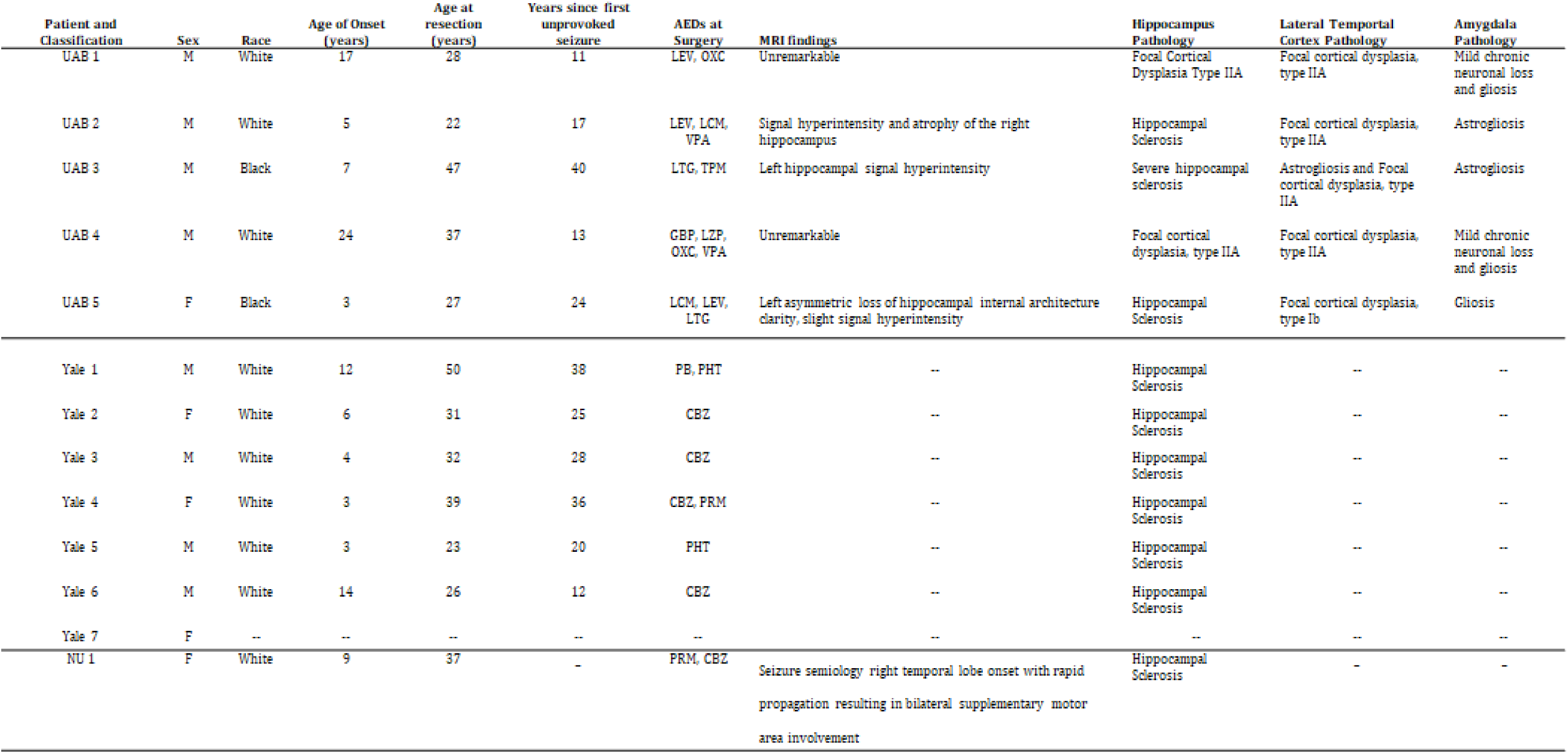
Human demographics from resected hippocampal tissue from TLE patients.

### 2.5 Kainate Treatment

Animals were injected with kainic acid (KA) [10 mg/kg; (Tocris Cookson Inc., Ellisville, MO, USA)] or saline (vehicle) intraperitoneally (IP). The severity of behavioral seizures following KA injection was scored according to the Racine scale [24]: a five-point scale which takes the five following behaviors as indicative of respectively increasing seizure severity: mouth and face clonus and head nodding (1); clonic jerks of one forelimb (2); bilateral forelimb clonus (3); forelimb clonus and rearing (4); forelimb clonus with rearing and falling (5). The onset of status epilepticus (SE) was defined as the time from KA injection to the occurrence of continuous seizure activity (Racine score 4-5) over a period of 4 hours. All control animals were handled in the same manner as the KA-treated animals but injected with saline. For tissue collection, the hippocampus was removed and oxygenated (95%/5% O2/CO2) in ice-cold cutting solution (110 mM sucrose, 60 mM NaCl, 3 mM KCl, 1.25 mM NaH2PO4, 28 mM NaHCO3, 0.5 mM CaCl2, 7 mM MgCl2, 5mM glucose, 0.6 mM ascorbate). The *cornu ammonis* (all CA regions), and the dentate gyrus (DG) region were microdissected and frozen immediately on dry ice. The hippocampus was bisected with the dorsomedial half being divided into four pieces. Using anatomic landmarks, each piece was dissected into CA and DG region. The CA and DG were dissected with a cut along the hippocampal fissure. The tissue was then stored at −80°C for RNA and DNA extraction.

### 2.6 Small Animal Magnetic Resonance Imaging

*T*_*1*_- and *T*_*2*_-weighted images were collected on a 9.4T Bruker BioSpin horizontal small bore animal MRI scanner. The imaging parameters were set as follows: 1 mm slice thickness, 1 mm between slice distance, 0.1 x 0.1 x 1 mm voxel size, 30 x 30 mm FOV, 27 images per acquisition. T_2_-weighted hippocampal intensities were normalized to within-slice cortical intensity using ImageJ software (n=5/group).

### 2.7 Western Blotting

Protein concentrations were estimated by Bradford Assay (Biorad), and 25µg of total protein/sample was reduced in 5x sample loading buffer (0.1 M Tris-HCl, 4% SDS, 20% glycerol, 0.2% β-mercaptoethanol, 0.2% bromphenol blue), boiled for 10 min, separated by 10% SDS-PAGE, and transferred onto PVDF membranes using Trans-Blot Turbo transfer system (1704155, BioRad, Hercules, CA, USA). Membranes were activated with methanol for three minutes before transfer, blocked for 1hr at room temperature and incubated overnight at 4°C with primary antibodies following the transfer. Three washes were done with 1x PBST (PBS and 0.01% Tween) between primary and secondary antibodies and after stripping. The membranes were blocked with 1:1 Licor Blocking buffer (P/N 927-40003, Licor) and PBST for one hour at room temperature after transfers and stripping. Imaging was done using Licor Odyssey scanner at 700/800 channel, and Licor Odyssey software. Image analysis was done using Image Studio Lite Ver. 3.1.

### 2.8 Sample Preparation for mass spectrometry

Protein was extracted from rat dorsal hippocampus CA using M-PER (78501, Thermo Fisher Scientific) and quantified using Pierce BCA Protein Assay Kit (23225, Thermo Fisher Scientific). Extracts were diluted in LDS PAGE buffer (NP0007, Invitrogen) followed by reduction, heat denaturing, and separation on an SDS Bis-Tris gel (4-12%, NP0323BOX, Invitrogen). The gels were stained overnight with colloidal blue (89871, Invitrogen). The entire lane comprising each sample was cut into 12 MW fractions and equilibrated in 100 mM ammonium bicarbonate (AmBc). Gel slices were reduced, carboxymethylated, dehydrated, and digested with Trypsin Gold (V5280, Promega, Madison, WI, USA) as per manufacturers’ instructions. Following digestion, peptides were extracted, the volume was then be reduced in a SpeedVac to near dryness, and resuspended to 20µl using 95% ddH_2_O/ 5% ACN/ 0.1% formic acid (FA) prior to analysis by 1D reverse phase LC-ESI-MS2 (as outlined below).

### 2.9 HPLC-electrospray tandem mass spectrometry

Peptide digests were injected onto a 1260 Infinity HPLC stack (Agilent, Santa Clara, CA, USA) and separated using a 75 micron I.D. x 15 cm pulled tip C-18 column (00G-4053-E0, Jupiter C-18 300 Å, 5 micron, Phenomenex, Torrance, CA, USA). This system runs in-line with a Thermo Orbitrap Velos Pro hybrid mass spectrometer, equipped with a nano-electrospray source (Thermo Fisher Scientific), and all data was collected in CID mode. The HPLC was configured with binary mobile phases that include solvent A (0.1%FA in ddH_2_O), and solvent B (0.1%FA in 15% ddH_2_O / 85% ACN), programmed as follows; 10min @ 0%B (2µL/ min, load), 120min @ 0%-40%B (0.5nL/ min, analyze), 15min @ 0%B (2µL/ min, equilibrate). Following each parent ion scan (350-1200m/z @60k resolution), fragmentation data (MS2) was collected on the topmost intense 15 ions. For data dependent scans, charge state screening and dynamic exclusion were enabled with a repeat count of 2, repeat duration of 15.0s, and exclusion duration of 60.0s.

### 2.10 Mass Spectrometry Data Conversion and Searches

The XCalibur RAW files were collected in profile mode, centroided and converted to MXML using ReAdW v. 3.5.1. The mgf files were then created using MzXML2Search (included in TPP v. 3.5) for all scans. The data was searched using SEQUEST, which was set for two maximum missed cleavages, a precursor mass window of 20ppm, trypsin digestion, variable modification C at 57.0293, and M at 15.9949. Searches were performed with a species-specific subset of the UniRef100 database.

### 2.11 Peptide Filtering, Grouping, and Quantification

The list of peptide IDs generated based on SEQUEST search results were filtered using Scaffold (Protein Sciences, Portland, OR, USA). Scaffold filters and groups all peptides to generate and retain only high confidence IDs while also generating normalized spectral counts (N-SC’s) across all samples for the purpose of relative quantification. The filter cut-off values were set with minimum peptide length of >5 AA’s, with no MH+1 charge states, with peptide probabilities of >80% C.I., and with the number of peptides per protein ≥2. The protein probabilities are then set to a >99.0% C.I., and an FDR<1.0. Scaffold incorporates the two most common methods for statistical validation of large proteome datasets, the false discovery rate (FDR) and protein probability [25-27]. Relative quantification across experiments were then performed via spectral counting, and when relevant, spectral count abundances were then normalized between samples [28-30].

### 2.12 Proteomics Analysis

For the proteomic data generated, two separate non-parametric statistical analyses are performed for each pair-wise comparison. These non-parametric analyses include 1) the calculation of weight values by significance analysis of microarray (SAM; cut off >|0.6|combined with 2) T-Test (single tail, unequal variance, cut off p < 0.05), which then were sorted according to the highest statistical relevance in each comparison. For SAM, whereby the weight value (W) is a statistically derived function that approaches significance as the distance between the means (μ1-μ2) for each group increases, and the SD (δ1-δ2) decreases using the formula, W=(μ1-μ2)/(δ1-δ2)[31, 32]. For protein abundance ratios determined with N-SC’s, we set a 1.5-2.0 fold change as the threshold for significance, determined empirically by analyzing the inner-quartile data from the control experiment indicated above using ln-ln plots, where Pierson’s correlation coefficient (R) was 0.98, and >99% of the normalized intensities fell between +/-1.5 fold. In each case, any two of the three tests (SAM, Ttest, or fold change) had to pass.

Gene ontology assignments and pathway analysis were carried out using MetaCore (GeneGO Inc., St. Joseph, MI, USA). In addition, the final proteins list is analyzed using the auto-expand algorithm within MetaCore using the default setting (i.e. expanded by 50 nodes). In parallel, the expand-by-one algorithm is used to identify connections to the neighboring proteins, known drug interactions, and any known correlation to a disease, or specific biological process. Interactions identified within MetaCore are manually correlated using full-text articles. Detailed algorithms have been described previously [33, 34].

### 2.13 Human Electrophysiology

The electrophysiological data obtained from slice studies were derived from patients with medically intractable epilepsy undergoing elective neurosurgical tissue resection for the removal of a sclerotic hippocampus. All patients gave their informed consent, before surgery, for the use of the resected brain tissue for scientific studies. This study was approved by the Newcastle and the North Tyneside 2 Local Research Ethics Committee (06/Q1003/51) (date of review 03/07/06) and had clinical governance approved by the Newcastle Upon Tyne Hospitals NHS Trust (CM/PB/3707).

### 2.14 In vitro human neocortex recordings

Briefly, human cortical samples were derived from material removed as part of the surgical treatment of medically intractable cortical epilepsy from the mesial temporal lobe regions with the written informed consent of the patients. Slices were prepared from these samples using methods as previously described [35-37]. The time between resection and slice preparation was <5 min. Extracellular recordings (DC–500 Hz) were conducted with ACSF-filled glass microelectrodes (2 MΩ) connected to an extracellular amplifier (EXT-10-2F, npi electronic GmbH, Tamm, Germany). Signals were digitized (5 kHz) and recorded on a computer and then extracellular field recordings were analyzed to detect events using a custom-written code in Matlab2015b (Mathworks, MA, USA).

### 2.15 siRNA Infusion

For electrophysiological studies, hippocampal slices were collected from 6-8 week old, male Sprague-Dawley rats. All rats had previously undergone stereotactic cranial infusion of siRNA according to previously described methods [38]. Briefly, animals were anesthetized by way of intraperitoneal injection of dexmedetomidine-ketamine and received bilateral infusions of Accell SMARTpool siRNAs (Thermo) targeting either OGT (#E-080125-00-05) or scrambled, negative controls (#D-001910-10-05) in the dorsal hippocampus using the following stereotaxic coordinates relative to bregma: A/P −3.6mm, M/L±1.7mm, D/V −3.6mm. Infusions were delivered at a constant rate of 0.1 uL per minute using a linear actuator for a total volume of 1 uL per side. Non-targeting, fluorescent Accell siRNA (#D-001960-01) were used to confirm targeted delivery of siRNA to the dorsal hippocampus. For all conditions, fresh stocks of siRNA (100 μM) were re-suspended in Accell siRNA resuspension buffer to a concentration of 4.5 μM immediately prior to surgery.

### 2.16 Electrophysiology

Following surgery, each rat was allowed five days of recovery time after which its brain was harvested and hippocampal slices were collected for further testing. High-frequency stimulation of the Schaffer collateral/commissural pathway (CA3-CA1) was conducted using four trains of 100 pulses at 100 Hz, spaced 60 seconds apart. The initial slope of the field excitatory postsynaptic potential (EPSP) was measured as an index of synaptic strength. %fEPSP slopes were averaged after 20 min of baseline recording. Electrophysiological data are reported as means ±SEM, where n represents the number of slices.

### 2.17 Statistical Analysis for Biochemistry studies

Data is expressed as mean ±S.E.M and compared by a Student-test and Man-Whitney. Shapiro-Wilk and Kolmogorov-Smirnov statistics were done to take into account any age, sex, race, and post-mortem interval information, none of the listed factors are contributing to our results for the OGT or O-GlcNAc protein levels for the human experiments. Statistically significant differences between groups were defined as p<0.05.

## 3 Results

### 3.1 Hippocampal O-GlcNAcylation and OGT activity is decreased in epileptic rats

KA induced epilepsy has been shown to alter a variety of PTMs in proteins of the hippocampus. Therefore we sought to quantify global O-GlcNAcylation levels in the hippocampus 8 weeks post-SE when the animals had become fully epileptic and experienced self-convulsive seizures **(Fig 1a).** Analysis of protein O-GlcNAcylation in the CA regions of the hippocampus revealed significant decreases of O-GlcNAcylation in epileptic animals compared to controls (t_(4)_=13.02, *p*=0.0002, t_(8)_=2.363, *p*=0.0457; **Fig 1b-c**). To investigate this decrease further, we measured OGT protein levels in the same region and observed a significant decreased in OGT protein levels in the epileptic rats when compared to controls(t_(4)_=13.02, *p*=0.0002, t_(8)_=2.363, *p*=0.0457; **Fig. 1d**). In light of these results, we wanted to understand whether loss of OGT contributed to neuronal hyper-excitability. Following siRNA-mediated knockdown of OGT and high-frequency stimulation of the Schaffer collateral/commissural pathway (**Supplemental Fig. 1a-c**), we detected a trend toward increasing percent fEPSP and fEPSP slope, findings which suggest an increased rate of neuronal firing with reduction of OGT. At the same time, no changes were detected in the paired-pulse facilitation between groups, indicating that any changes in neuronal firing were due to changes in the postsynaptic neuron. Taken together, these results indicate a reduction of O-GlcNAc and OGT protein levels in the epileptic hippocampus and suggest a correlation between epilepsy and protein O-GlcNAcylation.

**Figure 1:**
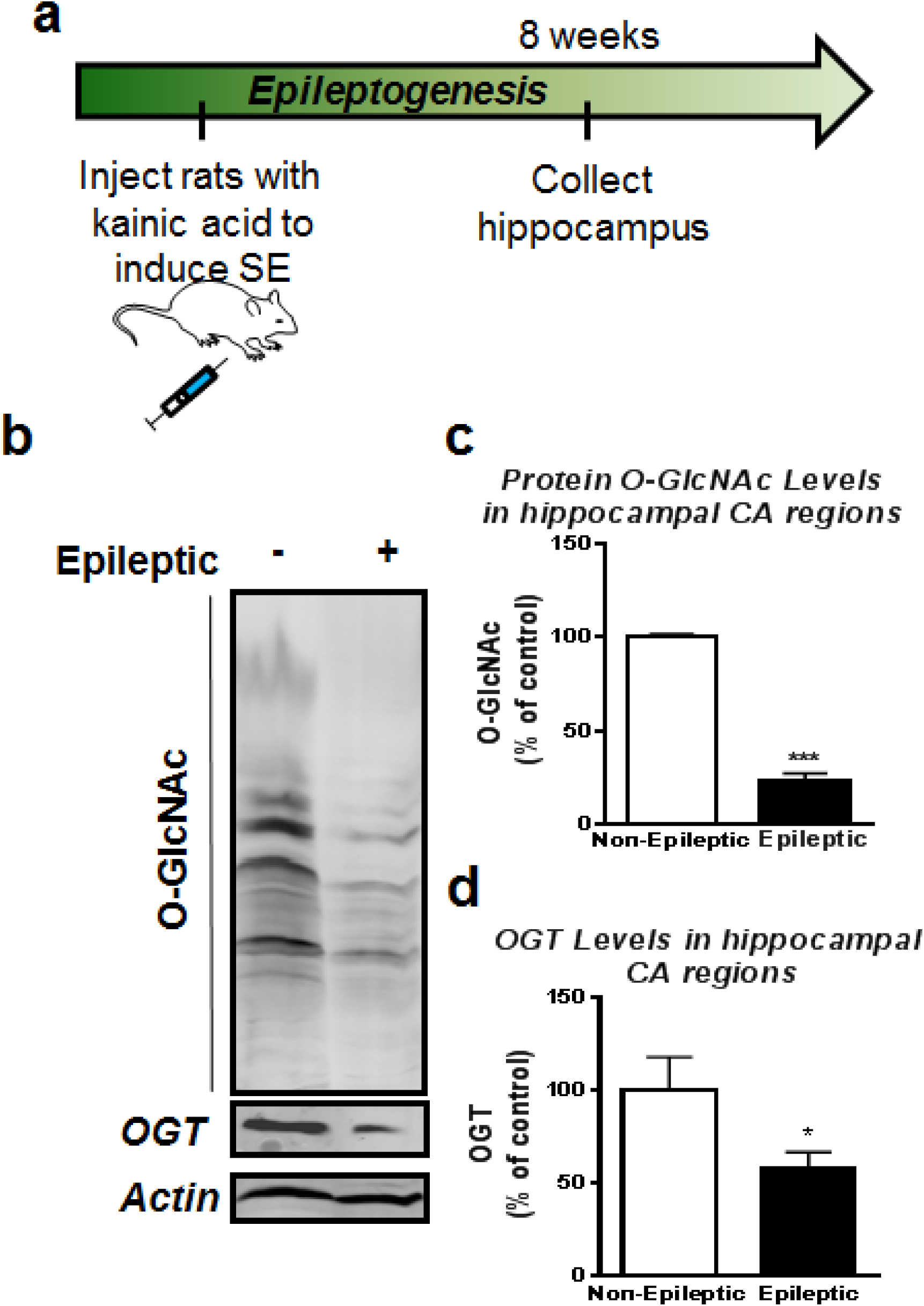
Hippocampus O-GlcNAcylation and OGT are decreased in epileptic rats. (a) Experimental design. Rats were either injected with saline or kainic acid in order to induce status epilepticus (SE). The animals were then sacrificed eight weeks later post kainic at which point these animals had become epileptic and the hippocampus was collected for protein analysis. (b) Representative O-GlcNAcylation as well as OGT and actin western blots for controls and epileptic rats. (c) Global O-GlcNAcylation was decreased in epileptic rats in comparison to control. (n=4-6 per group) (d) OGT protein levels were significantly reduced in epilepsy (n=4-6 per group). * denotes P <0.05 from controls, *** denotes P<0.001 from controls. Unpaired T-Test Error bars are SEM

### 3.2 Global protein O-GlcNAcylation changes in epileptic rats

Next, we sought to investigate specific proteins that displayed differential O-GlcNAcylation expression associated with TLE pathology. Using HPLC-electrospray tandem mass spectrometry we measured the abundance of proteins that had significantly altered O-GlcNAcylation in the CA regions of the hippocampus at 8 weeks post-SE. We found that 59 proteins were significantly differentially expressed in TLE, with seventeen of these 59 proteins exhibiting changes in O-GlcNAc marks. Gene ontology analysis revealed that the majority of diseases associated with differential expression of these proteins were neurodegenerative or cytoskeletal in nature. Additionally, among these seventeen proteins, twelve had been reported to be associated with epilepsy in the literature[3] (**Table1**) (**Fig 2a**).

**Figure 2:**
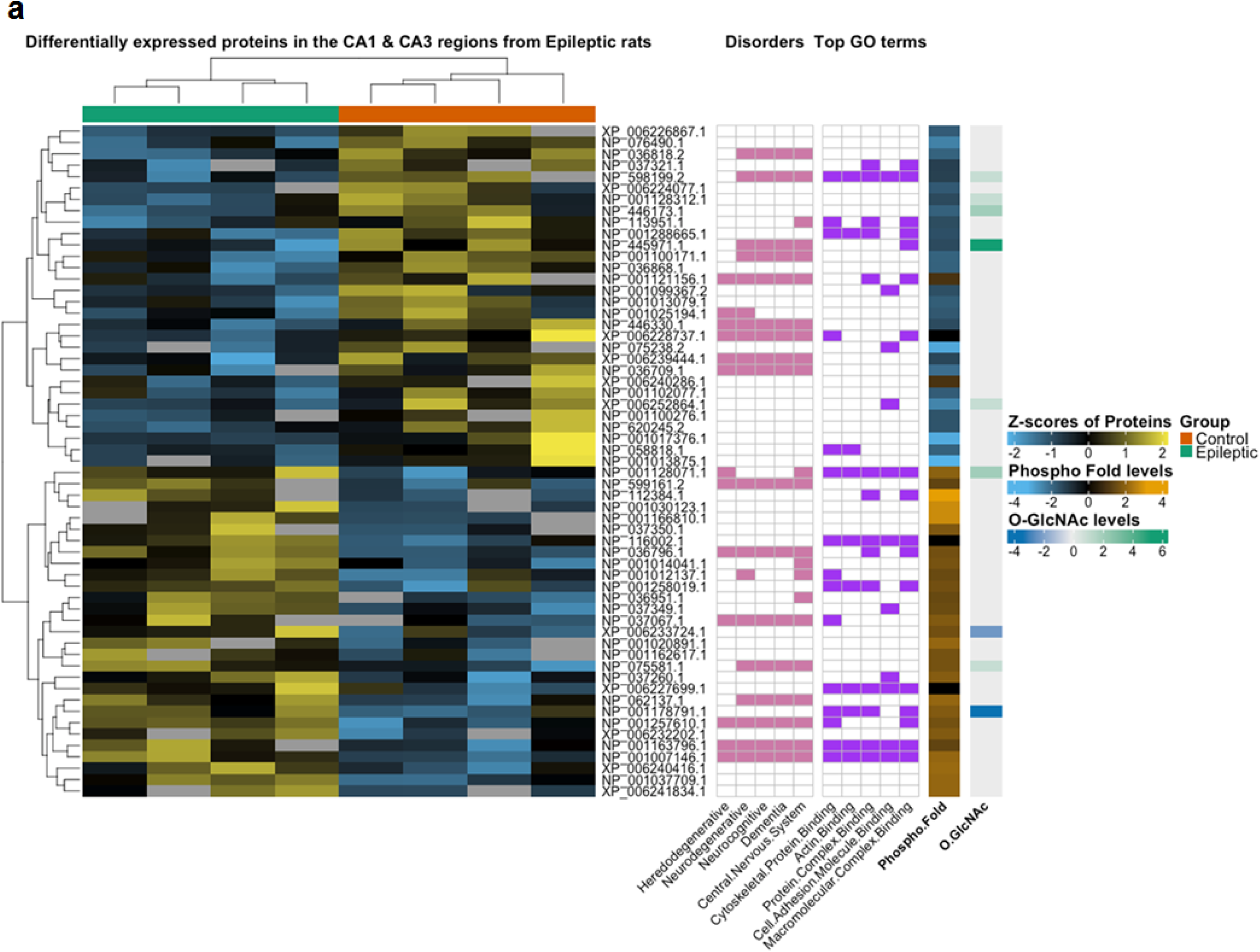
Global protein O-GlcNAcylation changes in Epileptic rats is protein dependent. (a) The heatmap illustrates all differentially expressed proteins (p<0.05) in epileptic rats (green bar) relative to controls (orange) bar. Each row is a protein indicated by the RefSeq accession number and each column in a biological replicate where the row and column order was determined by the Euclidian clustering method shown by the dendrograms. The protein values are shown as standardized z-scores, where the color indicates the standard deviation increasing (yellow) or decreasing (blue) relative to the mean (black). Grey blocks indicate missing values for the respective biological replicate. Further, for each protein, the top five disorders and GO terms (adjusted p-value<0.05) are annotated in pink and purple respectively. Lastly, the phosphorylation (phospho) fold change and O-GlcNAc levels are indicated for each differentially expressed protein.

In addition to measuring O-GlcNAcylation, we also measured phosphorylation and discovered increases in protein phosphorylation, particularly on those proteins that had shown decreases in protein O-GlcNAcylation. We next analyzed overall protein expression against protein phosphorylation and protein O-GlcNAcylation (**Supplemental Fig.2a-b**) revealing two distinct cluster groups. These clusters indicated that increased protein expression was positively correlated with increased protein phosphorylation, and only a few of the more highly expressed proteins also had changes in O-GlcNAcylation. In contrast to phosphorylation, increased protein O-GlcNAcylation was predominantly seen in proteins with decreased in expression. These clusters persisted when O-GlcNAcylation and phosphorylation were plotted against each other (**Supplemental Fig.2c).** The Z-scores were plotted from each biological replicate against either modification to demonstrate the contrast between their fold change **(Supplemental Fig.2d-e)**. Taken together our mass spectrometry analysis corroborated our findings that overall protein O-GlcNAcylation was decreased in the epileptic animal hippocampus while highlighting the particular proteomic ontologies affected by this loss. Additionally, our findings revealed that certain proteins actually show increased O-GlcNAcylation in the epileptic hippocampus. Collectively, these findings provide evidence that differentially expressed proteins and changes in PTMs are associated with TLE and other disease states highlighting the importance of protein PTM in homeostasis.

### 3.3 OGA inhibition in the epileptic hippocampus via acute Thiamet-G treatment reduces epileptiform activity

The observed global loss of O-GlcNAcylation and OGT prompted additional experiments to determine the role of this PTM in epilepsy. Using the KA model of epilepsy, we recorded cortical brain activity and seizures with EEG one month post-SE. We then administered Thiamet-G (10mg/kg/day), a known OGA inhibitor used to increase O-GlcNAcylation, once a day for three consecutive days in order to measure its effect on epileptiform brain activity (**Fig 3a**). We measured baseline EEG activity between control animals and epileptic animals and found that epileptic animals demonstrating higher power than the controls indicating more epileptiform activity (**Fig 3b-c**). The epileptic rats presented with sharp spikes and larger amplitudes than the control animals that depicted synchronous activity or seizures in the spectrogram with warmer colors. These epileptic animals then underwent a daily regimen of OGA inhibition for three days while having their brain activity measured (**Fig 3-d**). Following three days of Thiamet-G treatment, epileptic rats displayed a reduction in epileptic waveform and a decrease in the number of seizures experience per day and seizure duration (t_(7)_=1.999,*p*=0.858, t_(34)_=3.497, *p*=0.0013; **Fig. 3e-f**). We then unraveled the spectrogram using a power spectrum in order to measure the significant changes in power between frequencies for each day of Thiamet-G treatment. (**Fig. 3g**). The frequencies were divided into bands of brainwaves that are characterized by their range in frequency and behavioral characteristics. For instance, lower band frequencies such as delta and theta waves are associated with sleep, while higher frequency bands such as gamma are more closely associated with consciousness and attentiveness [39]. These bands can be used to characterize seizure severity. In this study, OGA inhibition helped restore the power of the lower frequencies (delta-alpha) more so than the higher frequencies (beta-gamma) to the baseline of the control group.

**Figure 3:**
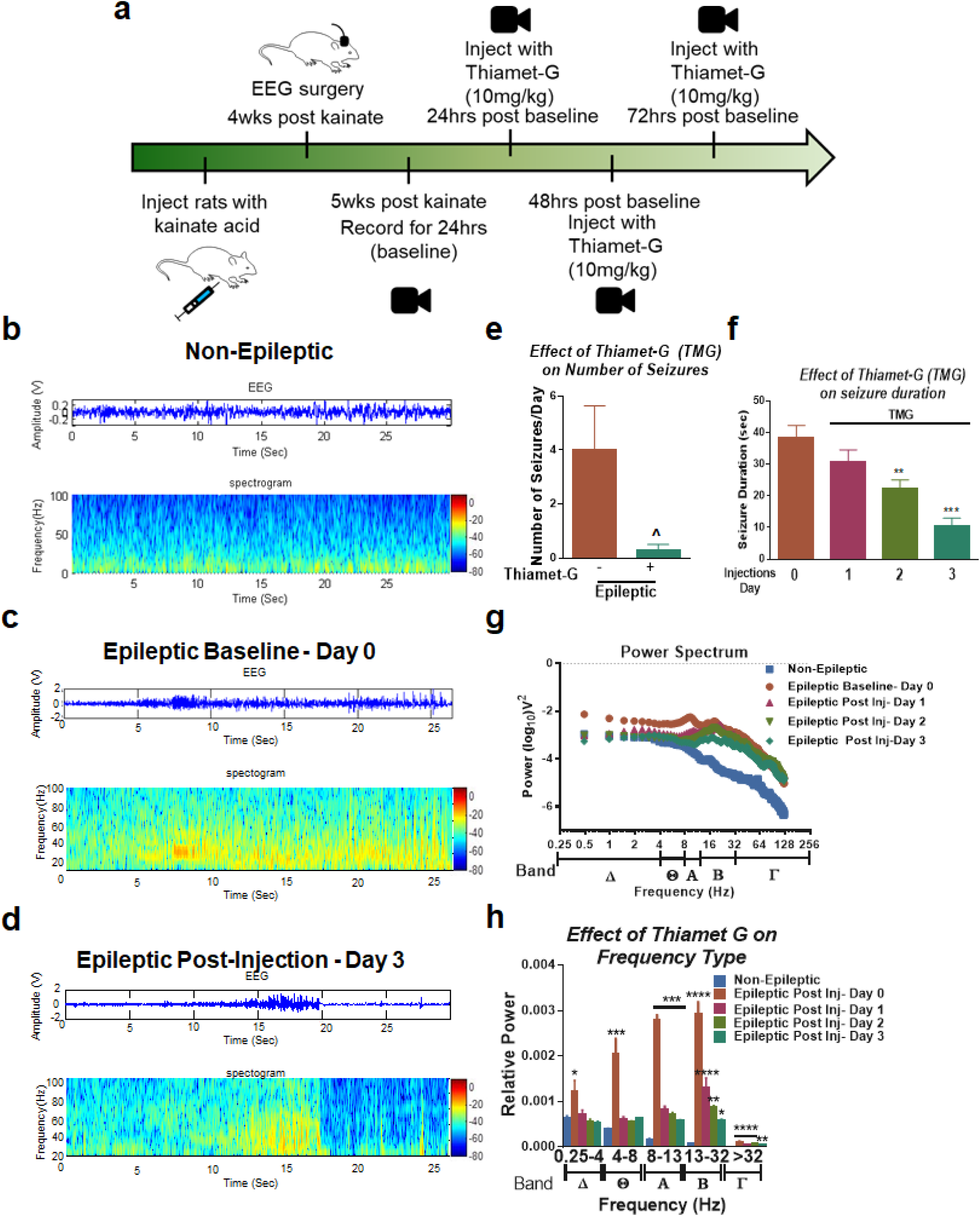
OGA inhibition decreases seizure duration and epileptiform activity. (a) Experimental outline. Epileptic rats were created using kainic acid. Four weeks post kainate the rats underwent EEG surgery where cortical electrodes were placed and the animals had a week to recover from the surgery before recordings were initiated. Baseline recordings were taking for 24hrs and Thiamet-G treatment ensued immediately after for three consecutive days followed by euthanization. (b) Cortical baseline EEG spectrogram of a saline (control) treated rat. (c) Cortical baseline EEG spectrogram of an epileptic rat during a seizure. (d) Cortical EEG spectrogram of the same epileptic rat following three days of Thiamet-G treatment. (e) The number of seizures decreased after three days of Thiamet-G treatment between the pre and post-treated animals. (f) Thiamet-G significantly decreased seizure duration by the second day of treatment and continued to decrease seizure duration up to the last day of treatment. (g) A power spectrum analysis demonstrated that the frequencies that were most dampened by Thiamet-G intervention were theta through gamma bands (h) Quantification of the power spectrum illustrates which frequencies were significantly decreased after treatment in comparison to control non-epileptic animals. * denotes P <0.05 from controls, ** denotes P<0.01 from controls, *** denotes P<0.001 from controls, **** denotes P<0.0001 from controls. ^ denotes P<0.10 One-way ANOVA, Error bars are SEM

We furthered analyzed each frequency type against their relative power. As expected the largest powers for each given frequency band were from the epileptic rat recordings prior to treatment (t(_32-52_)=, p=0.0016-<0.0001; **Fig 3h**). By the first day of treatment, these bands showed a reduction in power and began to mirror the power levels of the non-epileptic rats, with the exception of the gamma frequency. This discrepancy between the gamma frequency and the trend from the other bands could be explained by a local measure of activity and not by an overall global cortical network due to a single measurement of activity with an electrode. With each day of treatment, the relative power of each band decreased with the exception of the theta band which plateaued immediately after the first treatment of Thiamet-G. This band is typically characterized by the excitatory regular spiking, and intrinsic bursting pyramidal neurons, suggesting that inhibition of OGA via Thiamet-G may preferentially target this group of neurons more readily.

### 3.4 Chronic inhibition of OGA activity in epileptic rats increases hippocampal atrophy

Although OGA inhibition dampened epileptiform activity and seizure duration in a wide spectrum of frequencies, we sought to determine if there were any morphological changes associated with the Thiamet-G treatment over a prolonged period of usage. Hippocampal scarring and/or gliosis is often observed in animal models of TLE as well as in humans, where it leads to hippocampal atrophy. Hippocampal atrophy in TLE patients is observed using MRI T_2_ weighted scans where the ventricles adjacent to the hippocampus expand due to a reduction of size in the hippocampus. We created epileptic rats as previously described, and scanned these animals in an MRI machine at eight weeks post-injection in order to record their ventricular volumes prior to treatment. We then began a two-week treatment regimen for these animals with either saline or Thiamet-G (10mg/kg/day) and measured their ventricular volumes after treatment (**Fig 4a**). Coronal T_2_ pre/post scans were taken of saline and Thiamet-G injected rodents (**Fig.4b**). Voxels were quantified and compared to non-epileptic with saline injections for their respected time points (pre or post) (One way ANOVA, F=10.05, *p*= 0.0002 **Fig. 4c**). Epileptic rats displayed significantly higher voxel area units prior to treatment compared to non-epileptic controls. Following two weeks of treatment, voxel area increased in both Thiamet-G and saline-treated animals with no significant differences between the two groups. These scans suggest that Thiamet-G does little to inhibit or slow the progression of ventricular expansion seen in epilepsy [40-43].

**Figure 4:**
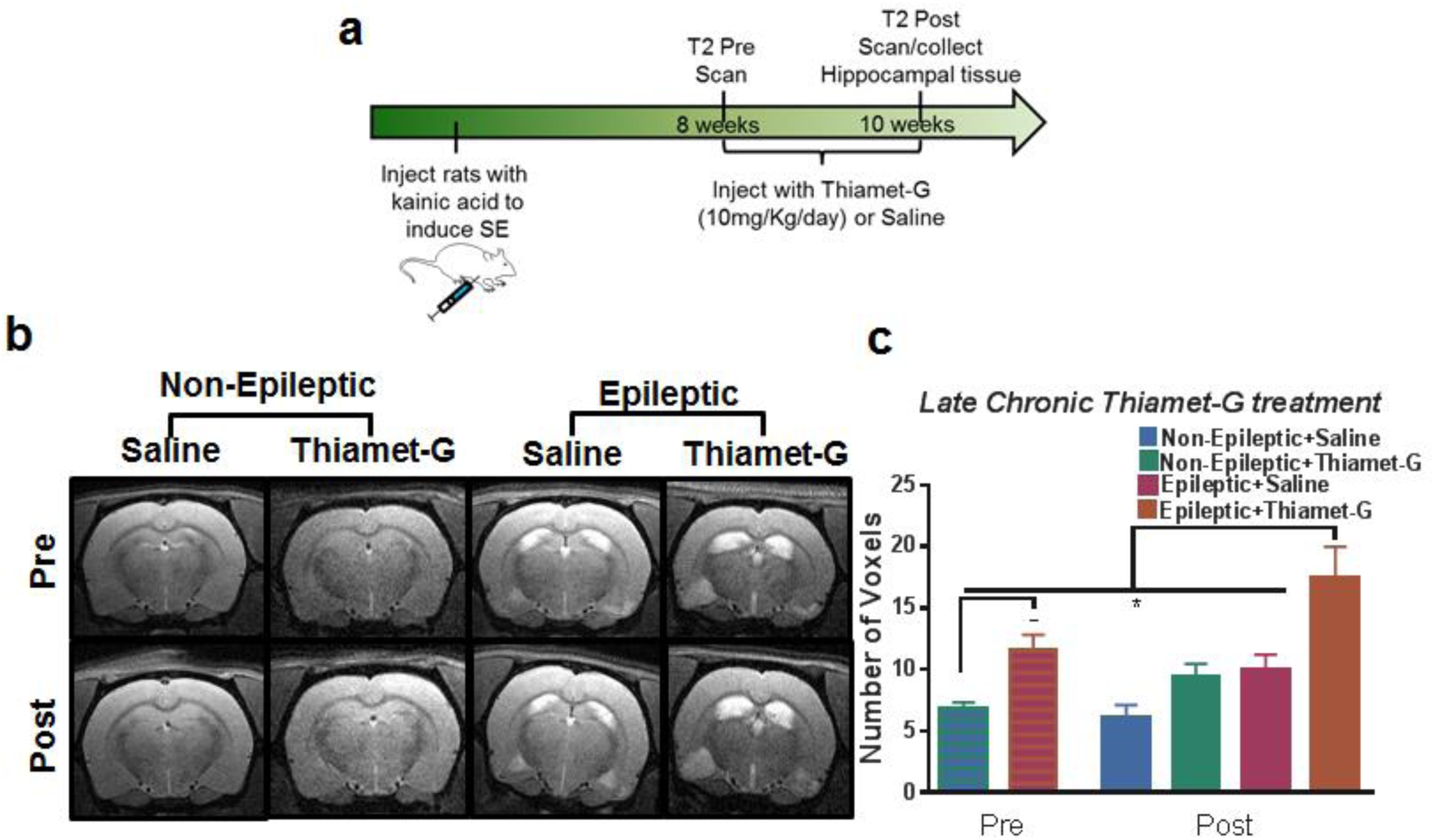
Thiamet-G treatment has no reduction in ventricle expansion and O-GlcNAcylation. (a) Experimental outline of animal model and treatment. Epileptic animals were created with kainic acid. Eight weeks post-kainate the animals had their first T2 scans were taken. Immediately following the scan, animals were treated with Thiamet-G (10mg/kg/day) for 2 weeks at the same time each day. The animals then had a final T2 scan where they were then sacrificed and the hippocampus was collected. (b)Representative pre/post T2 weighted images of epileptic and non-epileptic rats that were treated with either saline or Thiamet-G for two weeks. The CSF is bright white in the T2 MRI images demonstrating ventricle expansion with epilepsy and a more severe expansion with Thiamet-G treatment. (c) Quantification of T2 MRI images showing significant ventricle sizes between controls and epileptics before Thiamet-G treatment. Ventricle sizes significantly differed between the epileptic Thiamet-G treated group and the rest of the other group’s post-treatment. (n=8/group), * and – denotes P<0.05 from controls. One-way ANOVA. Error bars are SEM

Following MRI scans, animals were sacrificed and brain tissue was processed for immunohistochemistry experiments. We stained brain slices for GFAP as a marker for gliosis (**Supplemental Fig3a**) and for O-GlcNAcylation (**Supplemental Fig3b**). We observed increases in O-GlcNAcylation with Thiamet-G treatment as expected. However, with regards to GFAP, Thiamet-G increased its protein expression in healthy control rats but seemed to reduce GFAP expression in epileptic animals. Taken together, these experiments suggest that OGA inhibition does not stop or reverse epileptic hippocampal atrophy, but it may increase reactive astrocyte levels. These findings leave open the possibility that Thiamet-G treatment may slow the progression of hippocampal atrophy if it is begun earlier in the disease course. However, OGA inhibition does not appear to restore atrophied tissue.

### 3.5 Chronic Thiamet-G treatment in epileptic animals differentially alters OGA protein expression and O-GlcNAc substrates

Seeing as Thiamet-G treatment resulted in increased O-GlcNAcylation and decreased GFAP expression in epileptic animals we next wanted to understand how chronic treatment with Thiamet-G would affect O-GlcNAcylation levels on proteins shown to be differentially expressed in TLE. Specifically, we wanted to ask whether OGA’s expression was altered in epilepsy, and if so, whether Thiamet-G treatment restored OGA expression to homeostatic levels. We first looked at OGA protein expression in our epileptic animals that were treated for two weeks with Thiamet-G, (One way ANOVA F=1.852 p=0.085 **Fig. 5a-b**). We noticed no significant changes in OGA protein expression in control animals treated with Thiamet-G. Although not significant, we did notice a trend in increased OGA protein expression in epileptic animals. When these animals were treated with Thiamet-G, levels of OGA expression resembled those of saline-treated controls.

Based on our proteomic analysis (**Fig.2a**), we identified Sortilin-Related Receptor (SORL1) and tropomodulin 2 (Tmod2) as proteins that undergo increased and decreased protein O-GlcNAcylation in TLE, respectively (**Supplemental Fig.4**). SORL1 is a receptor that binds to LDL and transports it into the cells via endocytosis, a process that is subject to inhibition upon binding to the receptor-associated protein (RAP) [44]. SORL1 has also been implicated in APP trafficking to and from the Golgi apparatus and in Alzheimer’s disease [45, 46]. Tmod2 is an actin-binding protein that stabilizes ADP-bound actin monomers onto actin filaments and is downregulated in epilepsy [47, 48]. To test the effect of Thiamet-G administration on these proteins’ PTMs we used immunoprecipitation followed by Western blot to interrogate the levels of O-GlcNAcylaion on these proteins specifically. We observed no differences in O-GlcNAcylation om immunoprecipitated SORL1, nor did we find any differences in association with OGT (**Fig 5c**). Immunoprecipitation of Tmod2 revealed slight increases in O-GlcNAcylation in animals treated with Thiamet-G, along with decreases of total O-GlcNAcylation in the inputs, or the raw unimmunoprecipitated samples (**Fig 5d**). Furthermore, no differences were observed in the degree of association between Tmod2 and OGT. These results suggest that Thiamet-G cannot restore the decreased levels of O-GlcNAcylation of SORL1 and Tmod2 specifically in epileptic rats, a finding which led us to ask whether these observations are similar in human, resected TLE samples and whether Thiamet-G might have a greater impact on human O-GlcNAcylation than it did on rats.

**Figure 5:**
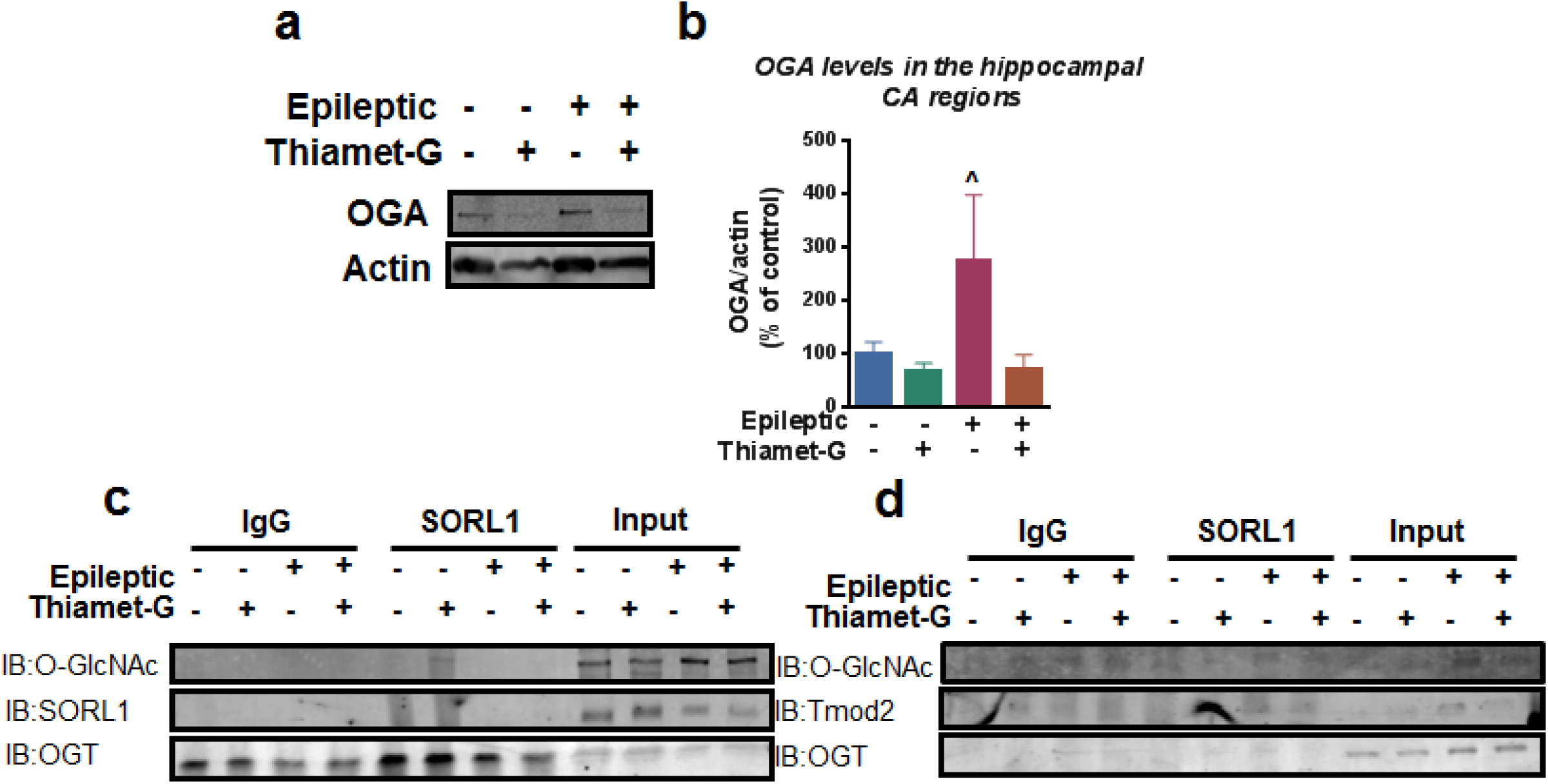
Thiamet-G treatment decreases OGA expression but leaves SORL1 and Tmod2 unmodified with O-GlcNAC. (a) Representative western blots of OGA and actin for the two-week saline or Thiamet-G treated epileptic and non-epileptic rats. (b) statistical analysis of the two-week saline or Thiamet-G treated epileptic and non-epileptic rats normalized to actin. (c) Immunoprecipitation of SORL1 with immunoblotting for O-GlcNAc (top membrane), SORL1 (middle membrane), and OGT (bottom membrane). (d) Immunoprecipitation of Tmod2 with immunoblotting for O-GlcNAc (top membrane), Tmod2 (middle membrane), and OGT bottom (membrane). (n=6-7/group) ^denotes P<0.10One-way ANOVA. Error bars are SEM.

### 3.6 Deficits in O-GlcNAcylation and OGT in patients with TLE

Our initial rodent studies have shown that O-GlcNAcylation and OGT are downregulated in the hippocampi of epileptic rats, but we were unsure of how O-GlcNAcylation and OGT might behave in human TLE tissue. We began by measuring O-GlcNAcylation and OGT expression in resected human hippocampus from TLE patients and compared them to age-matched controls from post-mortem human hippocampus tissue (**Fig. 6a**). We observed a significant loss of O-GlcNAcylation and OGT expression (t_(18)_=3.198, *p*=0.0050, t_(11)_=1.941, *p*=0.0783 **Fig 6b-c**) in TLE patients in comparison to postmortem tissue as seen in our epileptic rats. After recapitulating this loss of OGT and global O-GlcNAcylation in human tissue, we next asked whether SORL1 and Tmod2 were being modified in the same manner as we had seen in our epileptic rats. We immunoprecipitated SORL1 and Tmod2 and, in the rats, we observed no differences in O-GlcNAcylation of the proteins, nor any differences in their interaction with OGT (**Fig 6d**).

**Figure 6:**
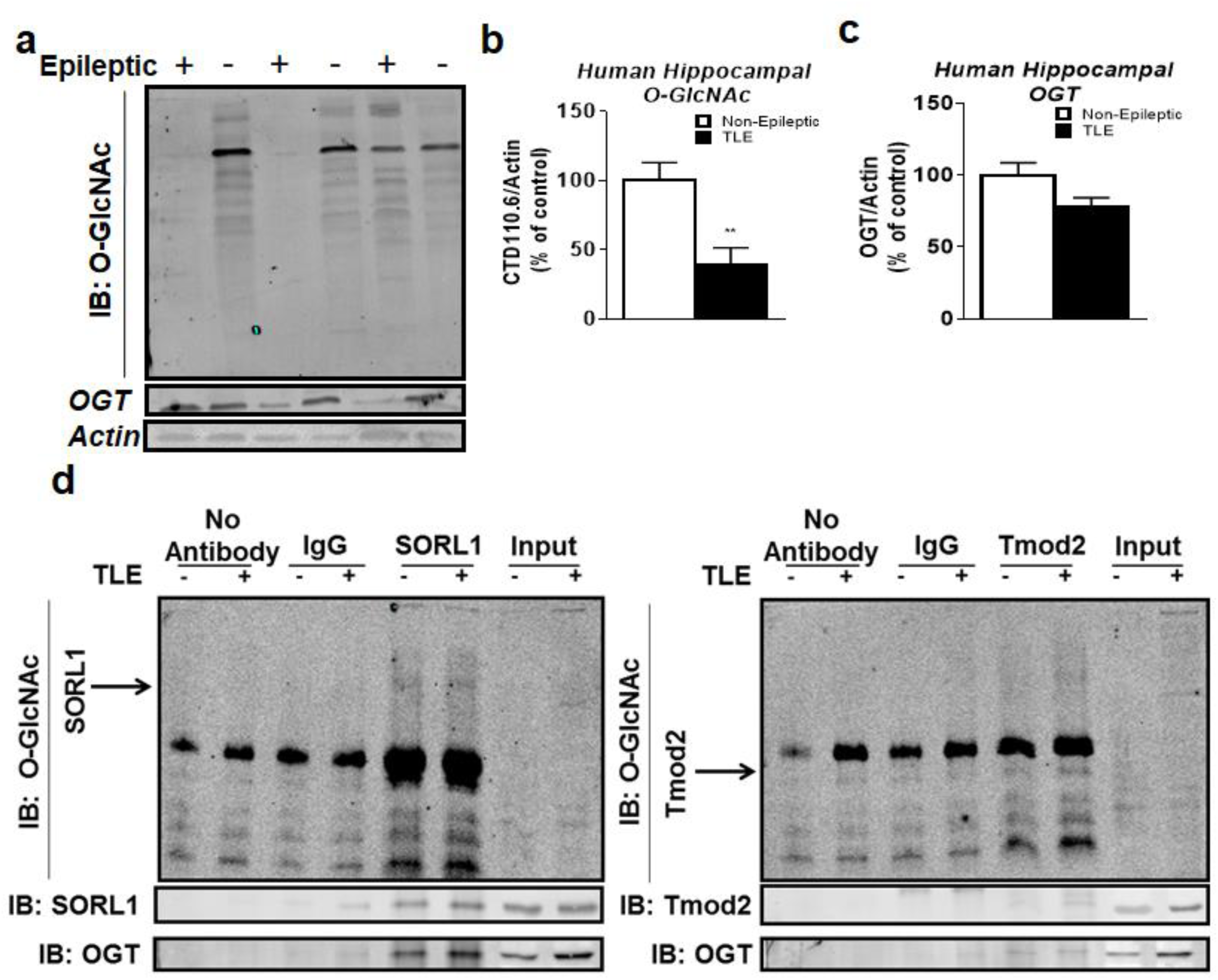
Deficits in O-GlcNAcylation and OGT in patients with TLE. (a) Western blot membrane with TLE human tissue and post-mortem non-epileptic alternating from left to right. The top membrane was probed with CTD110.6 antibody to show O-GlcNAc levels between both groups. The middle membrane was stripped and probed with OGT and the bottom membrane represents the level of actin between both groups. (b) Desensitization of O-GlcNAc levels between control and TLE individuals were quantified and actin was used to normalize O-GlcNAc. (n=11-13 per group) (c) Desensitization of OGT levels between control and TLE where taken and normalized to actin. (n=11-13 per group). (d) Immunoprecipitation of SORL1 and Tmod2 on resected TLE patients and postmortem tissue. Immunoblotting was performed with O-GlcNAc (top membrane), SORL1 (middle membrane), Tmod2 (middle membrane), and OGT (bottom membrane). Unpaired T-Test. ** denotes *P*<0.01 ^ denotes P<0.10 Error bars are SEM.

### 3.7 OGA inhibition in human TLE resected tissue decreases spike events and increases O-GlcNAcylation

Seeing as human TLE patient samples exhibited decreases in O-GlcNAcylation, we then wanted to know if restoring O-GlcNAcylation with Thiamet-G would decrease seizure spike events as observed in the rats treated with Thiamet-G. To test this hypothesis we obtained samples from TLE patients undergoing surgical resection. We immediately placed these samples in oxygenated ACSF, sectioned the tissue, and allowed them to acclimate to the bath for 1hr (**Fig 7a**). Following a 1hr incubation at room temperature, we recorded baseline activity for 1hr, finding that each slice exhibited spontaneous interictal-like activity. After 1 hr, slices were bathed in Thiamet-G (100 µM) for 3 hrs and the samples frozen for later molecular processing.

**Figure 7:**
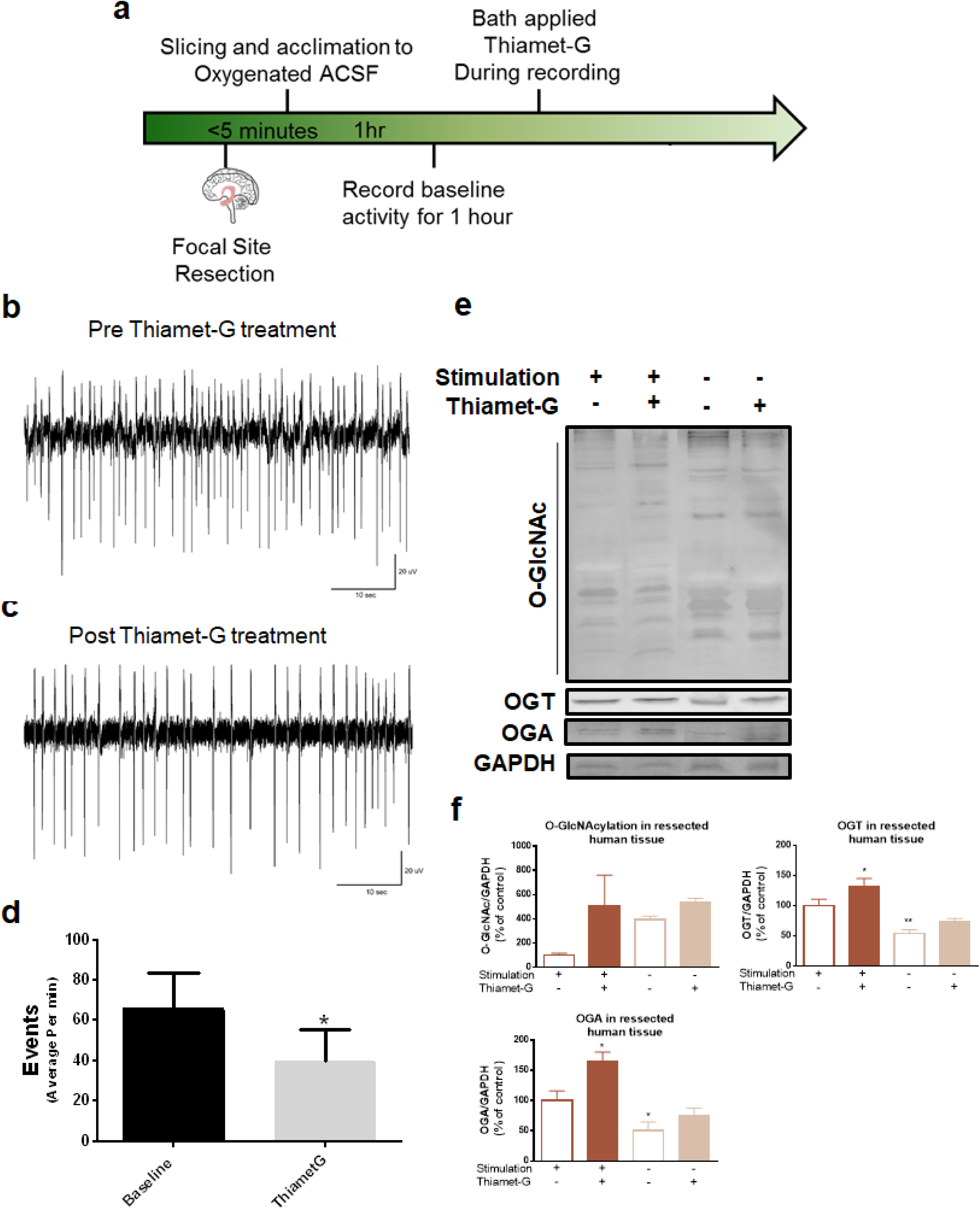
Thiamet-G bath application on human resected hippocampal focal site reduces seizures and decreases OGA protein expression and increases O-GlcNAcylation. (a) Experimental outlined. Samples were taken from patients that had gone temporal lobectomy that were unresponsive to AED. Samples were immediately placed in oxygenated ACSF and allowed to acclimate for 1hr. Baseline recording of activity was taken for 1hr followed by bath application of Thiamet-G. Samples were flash frozen and stored at −80°C. (b) Representative spike recording of resected hippocampus before Thiamet-G administration (c) Representative spike recording of the resected hippocampus after Thiamet-G bath application. (d) Quantification of spiking events per min of tissue slices at baseline and Thiamet-G administration. (e) Protein O-GlcNAcylation OGT and OGA were measured using western blots in order to ascertain Thiamet-G effects on the samples. (f) Quantification of western blot. O-GlcNAc and OGA were normalized to actin and compared to stimulated untreated group (n=4/group). * denotes P<0.05. Fisher LSD test. Error bars are SEM.

Prior to treatment with Thiamet-G, these slices exhibited an average of 66 spikes per minute, a rate which decreased to an average of 40 spikes per minute after one hour of treatment in Thiamet-G solution (**Fig7b-d**). Importantly, slices that were not treated with Thiamet-G showed no change in average spikes per minute over time. In this way, we showed that bath application of Thiamet-G to hyper-excitable human hippocampus significantly decreased spike frequency, recapitulating the similar effect that we observed *in vivo* in our epileptic rats.

We next examined the O-GlcNAcylation, OGT, and OGA levels in these tissues (**Fig 7e**). Protein O-GlcNAcylation generally increased with Thiamet-G treatment and decreased with vehicle control treatment depending on whether the tissue had electrophysiological recordings performed. (**Fig 7f**). OGT levels increased with recording and Thiamet-G treatment but not with Thiamet-G treatment alone; OGA, which is enzymatically inhibited by Thimaet-G increased during our electrophysiology recordings but showed little change with Thiamet-G treatment alone. Overall, we find that O-GlcNAcylation and OGT levels are decreased in epilepsy, but promoting this PTM pharmacologically resulted in decreased seizure frequency and, spikes, as well as increased protein O-GlcNAcylation.

## 4 Discussion

The current study demonstrates that O-GlcNAcylation and OGT are decreased in the CA1/CA3 regions of the hippocampus in a rodent model of epilepsy. By pharmacologically targeting this modification through pharmacological inhibition of OGA, we were able to raise global protein

O-GlcNAcylation levels, not only in epileptic rats but in resected human hippocampus tissue as well. To our knowledge, this is the first demonstration of a role for protein O-GlcNAcylation and its mediators in epilepsy. This is further supported by prior studies suggesting that this PTM is involved in prolonged seizure activity [3, 4]. Granted that O-GlcNAcylation is critical in modulating cellular homeostasis, we submit that this PTM shows promise as a new therapeutic target in epilepsy or TLE and other chronic seizures disorders [8-10]. Indeed, O-GlcNAc signaling has been characterized in numerous pathologies outside of the nervous system [49-52]. To date, O-GlcNAc has been limited to studies in the nervous system only with respects to Alzheimer’s disease, Parkinson’s disease, Huntington’s disease, schizophrenia, seizures, appetite, and synaptic plasticity [14-20, 53, 54]

Additionally, we demonstrated that the loss of OGT and O-GlcNAcylation does not present a homogeneous expression profile. We showed that although the majority of proteins showed decreased O-GlcNAcyaltion levels, there were some proteins that showed an increased in this PTM. Moreover, we demonstrated that proteins that have been associated with epilepsy had differentially expressed O-GlcNAc levels, changes that potentially alter their structure and function in TLE. Finally, we identified a novel biological target, OGA, which can be successfully depressed by Thiamet-G to promote O-GlcNAcylation levels and decrease the number of seizures and spikes *in vivo* both in rats and in human tissue. Although chronic inhibition of OGA in epileptic rats did not prevent or reverse ventricular expansion we did find that Thiamet-G treatment in epileptic animals and humans tissue could be used to reduce seizures and spike frequency. In future studies, it may be of interest to treat these rodents with Thiamet-G during earlier stages of epilepsy pathogenesis, during the onset of status epilepticus to investigate whether Thiamet-G can delay or halt epileptogenesis as a preventative treatment.

In summary, our results suggest that protein O-GlcNAcylation and its mediators play a previously unknown role in TLE and its animal models (**Fig 8**). These findings shed new light on the disorder and recommend novel therapeutic targets that warrant further study. Seeing as protein O-GlcNAcylation is closely tied to broader cellular metabolism, a program of treatment recognizes O-GlcNAcylation’s role in epileptic pathophysiology could employ many potential therapies targeting related pathways. These therapeutic candidates range from glucosamine to metformin and even the ketogenic diet, a therapy that would have fewer side effects compared to conventional AED’s [21, 55, 56].

**Figure 8:**
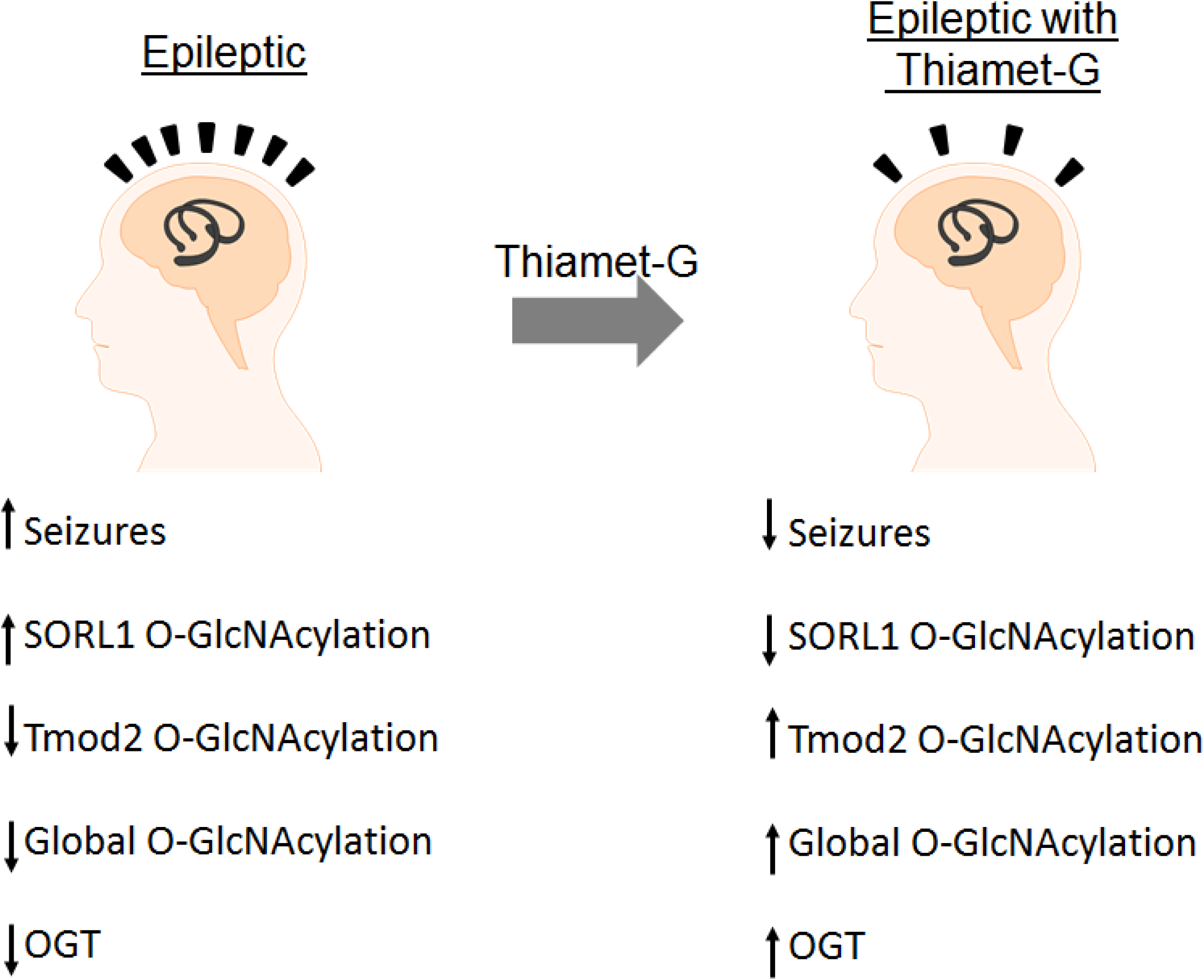
Protein O-GlcNAcylation in the epileptic hippocampus. We observed global losses of O-GlcNAcylation in human and rat TLE. However analyzing specific proteins and their modifications there were few that had increases namely SORL1, with the majority have losses of O-GlcNAc. In addition, we observe loss of OGT protein expression with epilepsy. By inhibition OGA we observed decreases in seizures and a restoration of protein O-GlcNAcylation homeostasis, in addition to increases in OGT.

## Supplemental Material

**Figure 1:**
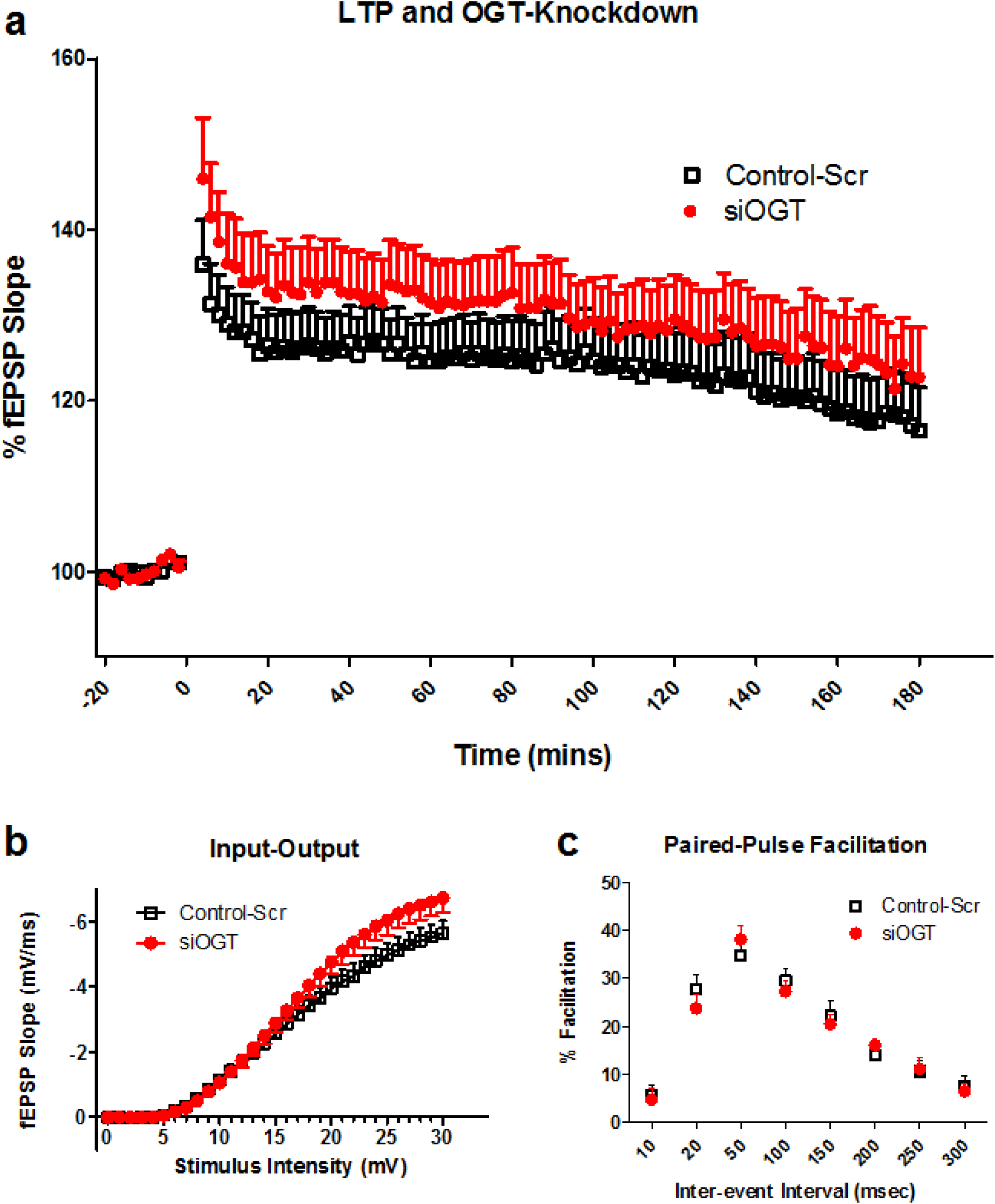
siRNA knockdown of OGT in the rat CA1 electrophysiological recordings. High-frequency stimulation of the Schaffer collateral/commissural pathway (CA3-CA1) was conducted using four trains of 100 pulses at 100Hz, spaced at 60 seconds apart. (a) The initial slope of the field excitatory postsynaptic potential (EPSP) as an index of synaptic strength. Recordings were taken for 180 minutes. (b) Percent fEPSP slopes were averaged after 20 minutes of baseline recording between input and output. (c) Paired-Pulse facilitation remained unchanged throughout the entire inter-event interval.

**Figure 2:**
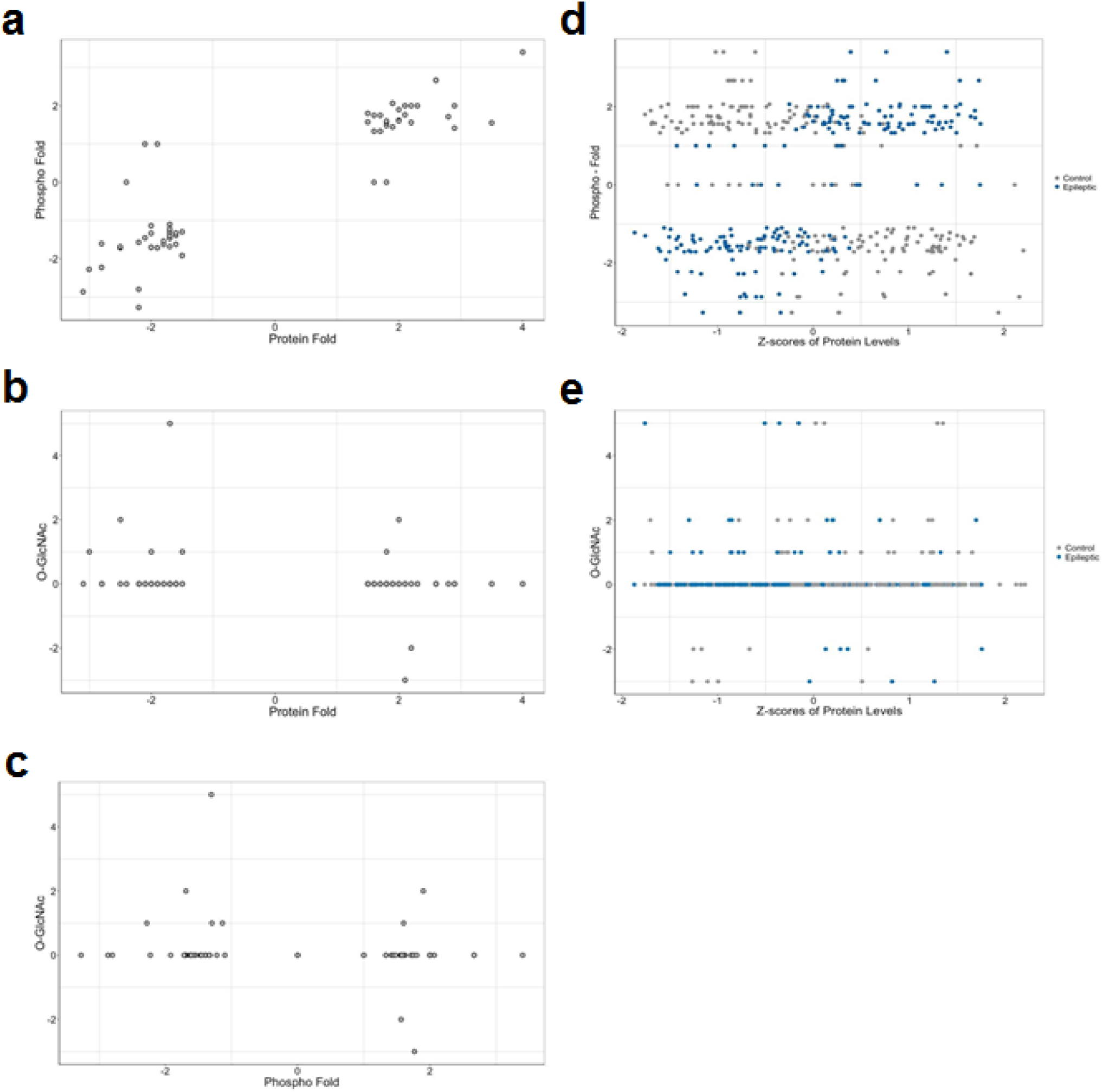
Relationship among protein expression, phosphorylation, and O-GlcNAc levels. For each differentially expressed protein, the phosphorylation and O-GlcNAc levels were shown in the scatter plots. The relationship is illustrated in (a) protein fold change and phosphorylation fold change (b) protein fold and O-GlcNAc levels and (c) phosphorylation fold and O-GlcNAc levels. To further demonstrate the relationship between protein levels and each modification, the z-scores from the proteins for each biological replicate are shown in contrast to (d) phosphorylation fold change and (e) O-GlcNAc levels.

**Figure 3:**
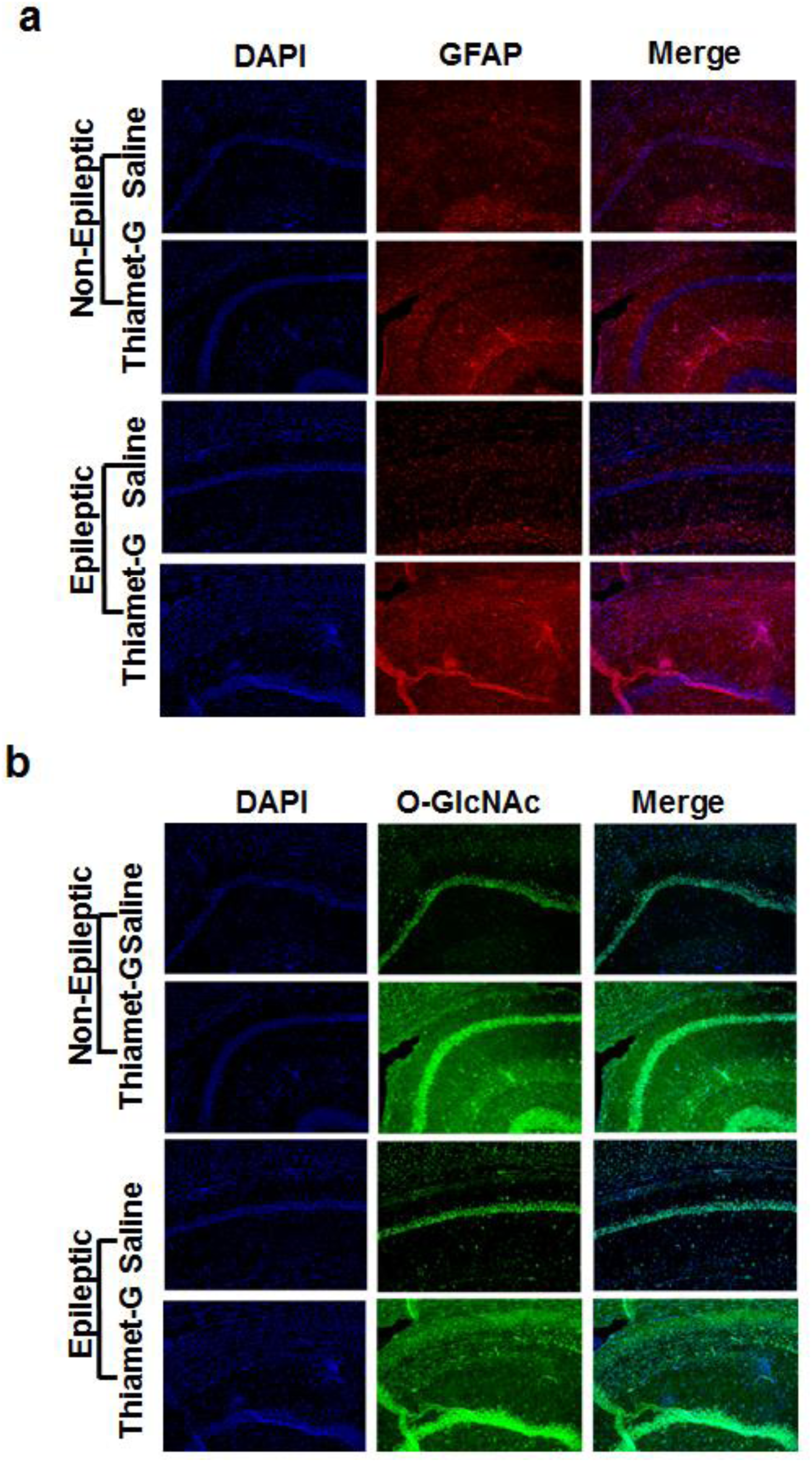
Immunohistochemistry staining of GFAP from two-week chronic Thiamet-G treatment on epileptic rats two-months post kainate. (a)Hippocampal CA region GFAP staining of two-month post kainate epileptic rats treated chronically with Thiamet-G or saline for two-weeks at 20x magnification (b) IHC staining of O-GlcNAcylation of the CA region of the hippocampus at 20x magnification of these animals post-Thiamet-G treatment illustrates increases in O-GlcNAcylation. (n=8/group).

**Figure 4:1.**
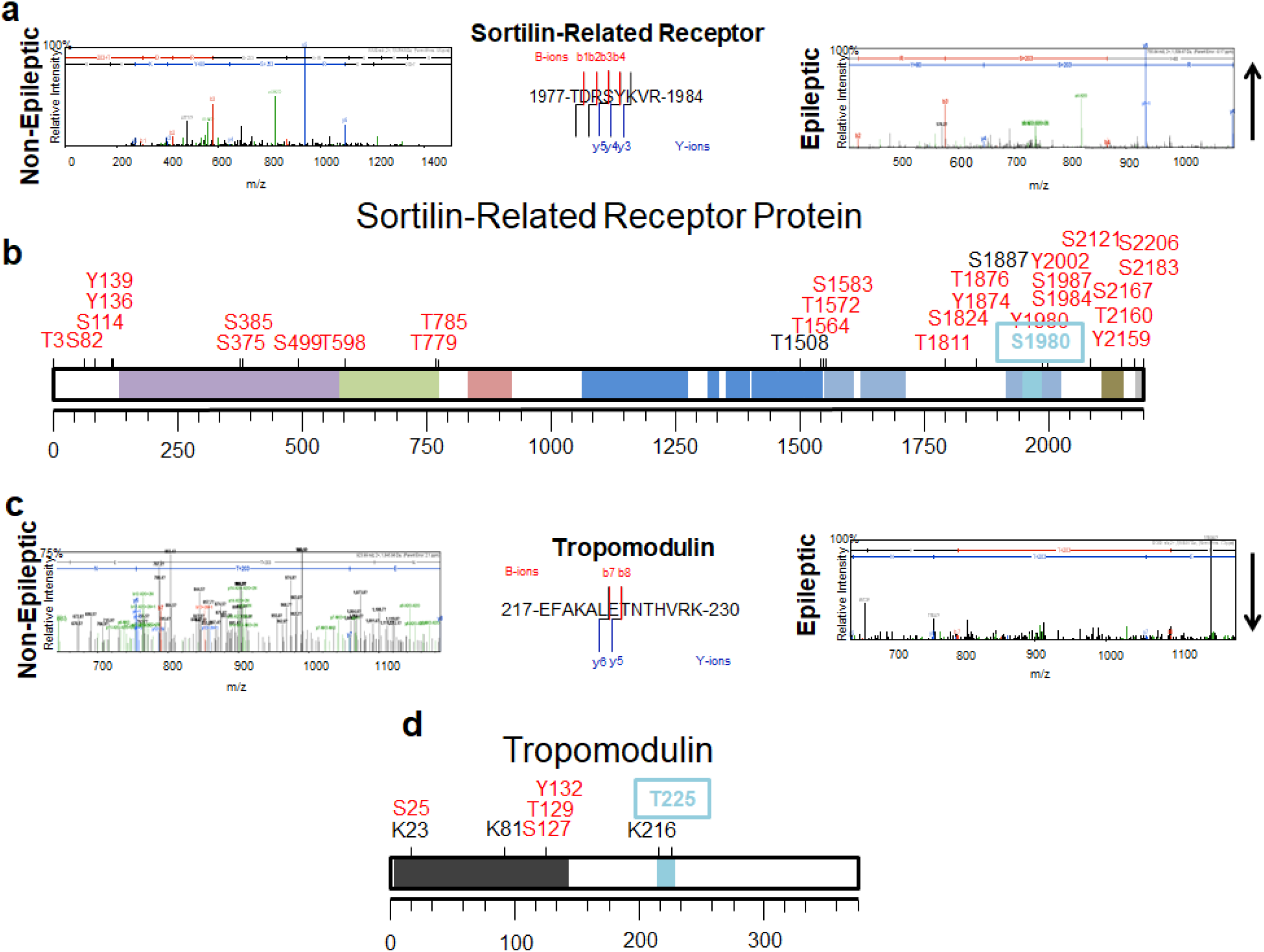
Sortilin-Related Receptor (SORL1) protein and Tropomodulin (Tmod2) Phosphorylated sites. (a) SORL1 peptide ^1977^TDRSYKVR^1984^ ms (m/z 755.85) for controls (left) and epileptic (right) CA hippocampus showed differentially O-GlcNAcylation on serine 1980. With an overall trend towards an increase in epilepsy. (b) SORL1 peptide; single letters represent the amino acid which has been cited as phosphorylated and the numbers represent the location of that amino acid on the protein. Colored bars represent domains: purple-sortilin-vsp10 domain, green-sortilin C domain, red-LDL receptor B domain, blue-LDL receptor A domain, light blue-Fibronectin type III domain, turquoise-peptide from our mass spectrometry, brown-transmembrane domain, grey-low complexity domain. (c) Tmod2 peptide ^217^EFAKALETNTHVRK^230^ MS (m/z 923.99) for controls (left) and epileptic (right) hippocampus that demonstrated overall decreases in O-GlcNAcylation at threonine 225 (d) Tmod2 peptide; single letters represent the amino acid which has been cited as phosphorylated and the numbers represent the location of that amino acid on the protein. Colored bars represent domains: black-tropomodulin domain, turquoise-peptide from our mass spectrometry.

## 6 Acknowledgments

We would like to thank Drs. Cristin Gavin and Jing Wang of the Evelyn F. McKnight Synaptic Plasticity Core at UAB for technical assistance with electrophysiology experiments. We would like also like to thank Dr. James Mobley for carrying out the mass spectrometry at University of Alabama at Birmingham (UAB) Mass Spectrometry/Proteomics Shared Facility. This work was supported by the Epilepsy Foundation, McNulty Civitan International Scientist Award, and the National Institute of Neurological Disorders and Stroke (NS090250).

## References

1. Liu, X.Y., et al., Comparative proteomics and correlated signaling network of rat hippocampus in the pilocarpine model of temporal lobe epilepsy. Proteomics, 2008. 8(3): p. 582–603.

2. Meriaux, C., et al., Human temporal lobe epilepsy analyses by tissue proteomics. Hippocampus, 2014. 24(6): p. 628–42.

3. Khidekel, N., et al., Probing the dynamics of O-GlcNAc glycosylation in the brain using quantitative proteomics. Nat Chem Biol, 2007. 3(6): p. 339–48.

4. Stewart, L.T., et al., Acute Increases in Protein O-GlcNAcylation Dampen Epileptiform Activity in Hippocampus. J Neurosci, 2017. 37(34): p. 8207–8215.

5. Gass, P., M. Kiessling, and H. Bading, Regionally selective stimulation of mitogen activated protein (MAP) kinase tyrosine phosphorylation after generalized seizures in the rat brain. Neurosci Lett, 1993. 162(1-2): p. 39–42.

6. Mielke, K., et al., Activity and expression of JNK1, p38 and ERK kinases, c-Jun N-terminal phosphorylation, and c-jun promoter binding in the adult rat brain following kainate-induced seizures. Neuroscience, 1999. 91(2): p. 471–83.

7. Nateri, A.S., et al., ERK activation causes epilepsy by stimulating NMDA receptor activity. Embo, 2007. 26(23): p. 4891–901.

8. Zachara, N.E. and G.W. Hart, Cell signaling, the essential role of O-GlcNAc! Biochimica et Biophysica Acta (BBA)-Molecular and Cell Biology of Lipids, 2006. 1761(5–6): p. 599–617.

9. Bond, M.R. and J.A. Hanover, A little sugar goes a long way: The cell biology of O-GlcNAc. The Journal of Cell Biology, 2015. 208(7): p. 869–880.

10. Hart, G.W., et al., Cross Talk Between O-GlcNAcylation and Phosphorylation: Roles in Signaling, Transcription, and Chronic Disease. Annual Review of Biochemistry, 2011. 80: p. 825–858.

11. Zachara, N.E., et al., Dynamic O-GlcNAc Modification of Nucleocytoplasmic Proteins in Response to Stress A SURVIVAL RESPONSE OF MAMMALIAN CELLS. Journal of Biological Chemistry, 2004. 279(29): p. 30133–30142.

12. Copeland, R.J., G. Han, and G.W. Hart, O-GlcNAcomics--Revealing roles of O-GlcNAcylation in disease mechanisms and development of potential diagnostics. Proteomics Clin Appl, 2013. 7(9-10): p. 597–606.

13. Yuzwa, S.A., et al., A potent mechanism-inspired O-GlcNAcase inhibitor that blocks phosphorylation of tau in vivo. Nat Chem Biol, 2008. 4(8): p. 483–90.

14. Gatta, E., et al., Evidence for an imbalance between tau O-GlcNAcylation and phosphorylation in the hippocampus of a mouse model of Alzheimer’s disease. Pharmacological research, 2016. 105: p. 186–197.

15. Wani, W.Y., et al., O-GlcNAcylation and neurodegeneration. Brain Res Bull, 2017. 133: p. 80–87.

16. Xie, S., et al., O‐GlcNAcylation of protein kinase A catalytic subunits enhances its activity: a mechanism linked to learning and memory deficits in Alzheimer’s disease. Aging cell, 2016.

17. Yuzwa, S.A., et al., Pharmacological inhibition of O-GlcNAcase (OGA) prevents cognitive decline and amyloid plaque formation in bigenic tau/APP mutant mice. Molecular Neurodegeneration, 2014. 9: p. 42.

18. Cole, R.N. and G.W. Hart, Cytosolic O-glycosylation is abundant in nerve terminals. Journal of Neurochemistry, 2001. 79(5): p. 1080–1089.

19. Lagerlof, O., et al., The nutrient sensor OGT in PVN neurons regulates feeding. Science, 2016. 351(6279): p. 1293–6.

20. Taylor, E.W., et al., O-GlcNAcylation of AMPA receptor GluA2 is associated with a novel form of long-term depression at hippocampal synapses. J Neurosci, 2014. 34(1): p. 10–21.

21. Cheung, W.D. and G.W. Hart, AMP-activated Protein Kinase and p38 MAPK Activate O-GlcNAcylation of Neuronal Proteins during Glucose Deprivation. Journal of Biological Chemistry, 2008. 283(19): p. 13009–13020.

22. Pekkurnaz, G., et al., Glucose regulates mitochondrial motility via Milton modification by O-GlcNAc transferase. Cell, 2014. 158(1): p. 54–68.

23. Lagerlof, O., G.W. Hart, and R.L. Huganir, O-GlcNAc transferase regulates excitatory synapse maturity. Proc Natl Acad Sci U S A, 2017. 114(7): p. 1684–1689.

24. Racine, R.J., Modification of seizure activity by electrical stimulation. I. After-discharge threshold. Electroencephalogr Clin Neurophysiol, 1972. 32(3): p. 269–79.

25. Weatherly, D.B., et al., A Heuristic method for assigning a false-discovery rate for protein identifications from Mascot database search results. Mol Cell Proteomics, 2005. 4(6): p. 762–72.

26. Keller, A., et al., Empirical statistical model to estimate the accuracy of peptide identifications made by MS/MS and database search. Anal Chem, 2002. 74(20): p. 5383–92.

27. <Nesvizhskii et al. -2003 -A Statistical Model for Identifying Proteins by Ta.pdf>.

28. Liu, H., R.G. Sadygov, and J.R. Yates, 3rd, A model for random sampling and estimation of relative protein abundance in shotgun proteomics. Anal Chem, 2004. 76(14): p. 4193–201.

29. Old, W.M., et al., Comparison of label-free methods for quantifying human proteins by shotgun proteomics. Mol Cell Proteomics, 2005. 4(10): p. 1487–502.

30. Beissbarth, T., et al., Statistical modeling of sequencing errors in SAGE libraries. Bioinformatics, 2004. 20 Suppl 1: p. i31–9.

31. Golub, T.R., et al., Molecular classification of cancer: class discovery and class prediction by gene expression monitoring. Science, 1999. 286(5439): p. 531–7.

32. Xu, W., et al., Serum profiling by mass spectrometry combined with bioinformatics for the biomarkers discovery in diffuse large B-cell lymphoma. Tumour Biol, 2015. 36(3): p. 2193–9.

33. Bhatia, V.N., et al., Software tool for researching annotations of proteins: open-source protein annotation software with data visualization. Anal Chem, 2009. 81(23): p. 9819–23.

34. Ekins, S., et al., Algorithms for network analysis in systems-ADME/Tox using the MetaCore and MetaDrug platforms. Xenobiotica, 2006. 36(10-11): p. 877–901.

35. Roopun, A.K., et al., A nonsynaptic mechanism underlying interictal discharges in human epileptic neocortex. Proc Natl Acad Sci U S A, 2010. 107(1): p. 338–43.

36. Cunningham, M.O., et al., Glissandi: transient fast electrocorticographic oscillations of steadily increasing frequency, explained by temporally increasing gap junction conductance. Epilepsia, 2012. 53(7): p. 1205–14.

37. Simon, A., et al., Gap junction networks can generate both ripple-like and fast ripple-like oscillations. Eur J Neurosci, 2014. 39(1): p. 46–60.

38. Webb, W.M., et al., Dynamic association of epigenetic H3K4me3 and DNA 5hmC marks in the dorsal hippocampus and anterior cingulate cortex following reactivation of a fear memory. Neurobiol Learn Mem, 2017. 142(Pt A): p. 66–78.

39. Cahn, B.R. and J. Polich, Meditation states and traits: EEG, ERP, and neuroimaging studies. Psychol Bull, 2006. 132(2): p. 180–211.

40. Coulter, D.A. and C. Steinhauser, Role of astrocytes in epilepsy. Cold Spring Harb Perspect Med, 2015. 5(3): p. a022434.

41. Wieshmann, U.C., et al., Development of hippocampal atrophy: a serial magnetic resonance imaging study in a patient who developed epilepsy after generalized status epilepticus. Epilepsia, 1997. 38(11): p. 1238–41.

42. Jackson, D.C., et al., Ventricular enlargement in new-onset pediatric epilepsies. Epilepsia, 2011. 52(12): p. 2225–32.

43. Dabbs, K., et al., Brain structure and aging in chronic temporal lobe epilepsy. Epilepsia, 2012. 53(6): p. 1033–43.

44. Bu, G., The roles of receptor-associated protein (RAP) as a molecular chaperone for members of the LDL receptor family. Int Rev Cytol, 2001. 209: p. 79–116.

45. Zollo, A., et al., Sortilin-Related Receptor Expression in Human Neural Stem Cells Derived from Alzheimer’s Disease Patients Carrying the APOE Epsilon 4 Allele. Neural Plast, 2017. 2017: p. 1892612.

46. Yin, R.H., J.T. Yu, and L. Tan, The Role of SORL1 in Alzheimer’s Disease. Mol Neurobiol, 2015. 51(3): p. 909–18.

47. Yang, J.W., et al., Aberrant expression of cytoskeleton proteins in hippocampus from patients with mesial temporal lobe epilepsy. Amino Acids, 2006. 30(4): p. 477–93.

48. Sussman, M.A., et al., Neural tropomodulin: developmental expression and effect of seizure activity. rain Res Dev Brain Res, 1994. 80(1-2): p. 45–53.

49. Macauley, M.S., et al., Elevation of Global O-GlcNAc in Rodents Using a Selective O-GlcNAcase Inhibitor Does Not Cause Insulin Resistance or Perturb Glucohomeostasis. Chemistry & Biology, 2010. 17(9): p. 949–958.

50. Champattanachai, V., R.B. Marchase, and J.C. Chatham, Glucosamine protects neonatal cardiomyocytes from ischemia-reperfusion injury via increased protein-associated O-GlcNAc. Am J Physiol Cell Physiol, 2007. 292(1): p. C178–87.

51. Singh, J.P., et al., O-GlcNAc signaling in cancer metabolism and epigenetics. Cancer Lett, 2015. 356(2 Pt A): p. 244–50.

52. Lewis, B.A. and J.A. Hanover, O-GlcNAc and the Epigenetic Regulation of Gene Expression.

53. Narayan, S., et al., Evidence for disruption of sphingolipid metabolism in schizophrenia. J Neurosci Res, 2009. 87(1): p. 278–88.

54. Hung, W.Y., D.E. Mold, and A. Tourian, Huntington’s-chorea fibroblasts. Cellular protein glycosylation. Biochem J, 1980. 190(3): p. 711–9.

55. Hart, G.W., Three Decades of Research on O-GlcNAcylation-A Major Nutrient Sensor That Regulates Signaling, Transcription and Cellular Metabolism. Frontiers in Endocrinology, 2014. 5: p. 183.

56. Chen, Y.-J., et al., Protective effects of glucosamine on oxidative-stress and ischemia/reperfusion-induced retinal injury. Investigative Ophthalmology & Visual Science, 2015. 56(3): p. 1506–1516.

